# Rescuing DNMT1 Fails to Fully Reverse the Molecular and Functional Repercussions of Its Loss in Mouse Embryonic Stem Cells

**DOI:** 10.1101/2024.05.07.592204

**Authors:** Elizabeth Elder, Anthony Lemieux, Lisa-Marie Legault, Maxime Caron, Virginie Bertrand-Lehouillier, Thomas Dupas, Noël J-M Raynal, Guillaume Bourque, Daniel Sinnett, Nicolas Gévry, Serge McGraw

## Abstract

Epigenetic mechanisms are crucial for developmental programming and can be disrupted by environmental stressors, increasing susceptibility to disease. This has sparked interest in therapies for restoring epigenetic balance, but it remains uncertain whether disordered epigenetic mechanisms can be fully corrected. Disruption of DNA methyltransferase 1 (DNMT1), responsible for DNA methylation maintenance, has particularly devastating biological consequences. Therefore, here we explored if rescuing DNMT1 activity is sufficient to reverse the effects of its loss utilizing mouse embryonic stem cells. However, only partial reversal could be achieved. Extensive changes in DNA methylation, histone modifications and gene expression were detected, along with transposable element de-repression and genomic instability. Reduction of cellular size, complexity and proliferation rate were observed, as well as lasting effects in germ layer lineages and embryoid bodies. Interestingly, by analyzing the impact on imprinted regions, we uncovered 20 regions exhibiting imprinted-like signatures. Notably, while many permanent effects persisted throughout *Dnmt1* inactivation and rescue, others arose from the rescue intervention. Lastly, rescuing DNMT1 after differentiation initiation worsened outcomes, reinforcing the need for early intervention. Our findings highlight the far-reaching functions of DNMT1 and provide valuable perspectives on the repercussions of epigenetic perturbations during early development and the challenges of rescue interventions.

**HIGHLIGHTS:**

- Extensive changes to epigenomic landscapes and gene expression following transient loss of DNMT1 activity
- Dysregulation of known imprinted regions and identification of 20 regions with imprinted-like signatures
- De-repression of MERVL and MT2 LTRs with evidence of chimeric gene transcript generation
- Shorter telomeres, DNA damage accumulation and reduction of cell size, internal complexity and proliferation rate
- Lasting effects upon differentiation toward germ layer lineages and embryoid bodies
- Worsened molecular and cellular outcomes when delaying *Dnmt1* rescue until after differentiation initiation

GRAPHICAL ABSTRACT

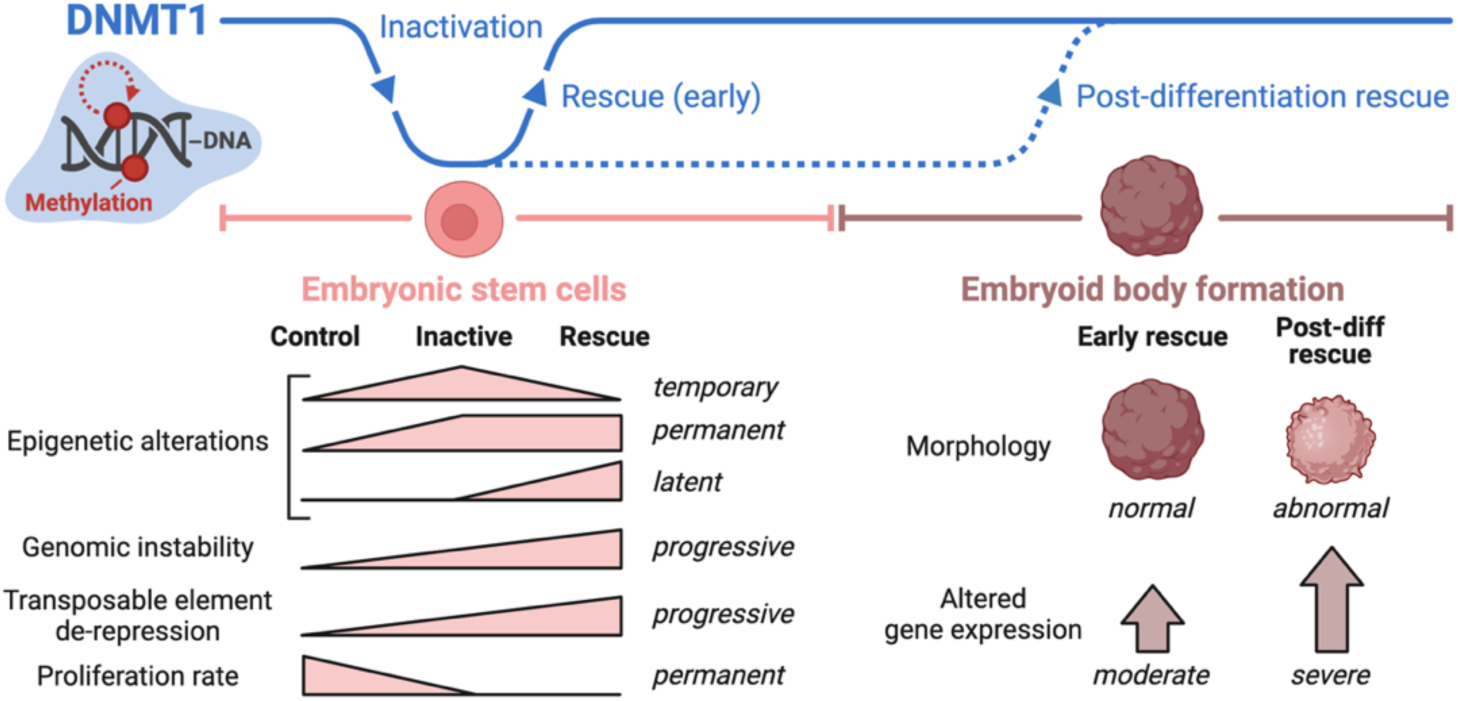

## INTRODUCTION

Epigenetic mechanisms, including DNA methylation and histone modifications, play a fundamental role in developmental programming by regulating molecular and cellular processes in response to internal and external stimuli without altering the underlying DNA sequence(1–4). Extensive research reveals that exposures to environmental stressors during development—such as toxins(5,6), poor nutrition(7–9), alcohol(10–12), assisted reproductive technology(13–17), and psychological stress(18,19)—can interfere with normal epigenetic programming, often with cell type(20–22) and sex(11,23) related specificities, thereby increasing susceptibility to various diseases and health issues. This has sparked a growing interest in therapeutic interventions aimed at mitigating epigenetic disruptions to reinstate molecular and cellular homeostasis(24–28). Yet, it remains uncertain if the detrimental impact of disordered epigenetic mechanisms can in fact be thoroughly remedied, all while avoiding potentially serious side effects.

DNA methyltransferase 1 (DNMT1) is a cornerstone of epigenetic regulation, primarily responsible for maintaining DNA methylation patterns during cellular division(29–32). DNA methylation maintenance by DNMT1 ensures epigenetic stability across cell generations(33), playing a critical role in genome integrity(34), genomic imprinting(35–37), transposable element silencing(38,39), gene expression regulation(40) and cell fate decisions(41–43). DNMT1 levels are finely tuned to specific developmental contexts; for instance, they are elevated in stem and progenitor cells and decrease during specialized differentiation(44–47). Aberrant DNMT1 activity—either deficiency or excess— disrupts DNA methylation profiles, leading to widespread hypomethylation(48–50) or localized hypermethylation(51–54), respectively. Beyond its catalytic function in DNA methylation, DNMT1 interacts with histone modifiers(55,56), chromatin remodelers(57) and non-coding RNAs(58), exerting influence over broader epigenetic networks(59,60). Given the wide-ranging scope of its mechanisms, dysregulation of DNMT1 has severe biological consequences, leading to embryonic lethality in mice(51,61) and contributing to the progression of various diseases such as developmental disorders(62), cancer(63,64), schizophrenia(65,66) and neurodegenerative pathologies(67–69). Emerging strategies for correcting abnormal DNA methylation profiles show promise for the treatment of disease(69–72). However, further research is needed to determine if DNA methylation perturbations can be rectified while fully reversing their effects on a broader molecular and cellular scale, without inflicting any collateral damage.

For mechanistic studies of epigenetic perturbations, embryonic stem cells (ESCs) serve as particularly useful models because of their pluripotency, rapid proliferation, relevance to both development and disease, and, importantly, their cellular plasticity(73–75). Cellular plasticity is a key characteristic that enables ESCs to tolerate substantial epigenetic alterations, allowing for stable cell cultures with aberrant expression of epigenetic enzymes while remaining in a pluripotent state and maintaining proliferative capacity(36,76–78). To investigate the feasibility of reversing DNA methylation perturbations, *Dnmt1*^tet/tet^ mouse ESCs (mESCs)(36) can be leveraged as a rescue model, wherein transient *Dnmt1* inactivation is achieved via the Tet-Off expression system(79), which facilitates reversible repression of an endogenous target gene with doxycycline treatment(79). Due to typical transactivation effects of the Tet-Off system(80), *Dnmt1*^tet/tet^ mESCs have higher DNMT1 levels relative to their wild-type counterpart; nevertheless, they function as a stable ES cell line(36,44,81) capable of differentiating along early developmental trajectories, with their derived embryoid bodies exhibiting comparable morphology to those from wild-type mESCs(44). The *Dnmt1*^tet/tet^ mESC model is therefore well-suited to explore whether restoring DNMT1 activity is sufficient for reversing the impact of its loss or if long-term effects persist post-rescue. Indeed, we and others have employed *Dnmt1*^tet/tet^ mESCs to demonstrate that loss of DNA methylation at imprinted regions is permanent when DNMT1-mediated DNA methylation maintenance is transiently interrupted(36,37), and we further showed that other, non-imprinted regions throughout the genome also become permanently hypomethylated(37).

Recognizing DNMT1’s far-reaching functions beyond DNA methylation maintenance, this study expands on our previous work to uncover the broader molecular and cellular alterations arising in *Dnmt1*^tet/tet^ mESCs after inactivating and rescuing *Dnmt1*. We profiled DNA methylation with enhanced coverage and sensitivity, five key histone modifications (H3K4me3, H3K27ac, H3K4me1, H3K27me3, H3K9me3), gene expression, transposable element silencing, genomic stability, as well as examined various cellular processes. Transient *Dnmt1* inactivation resulted in extensive epigenomic and gene expression alterations, de-repression of transposable elements and genomic instability. Reduction of cell size, internal complexity and proliferation rate were also detected, as well as lasting ramifications upon differentiation toward endoderm, mesoderm and ectoderm germ layers and embryoid bodies. Moreover, analyzing the impact on known imprinted regions enabled the identification of 20 regions exhibiting imprinted-like epigenetic and regulatory signatures. Notably, while many permanent effects were observed throughout *Dnmt1* inactivation and rescue, others were triggered by the rescue intervention itself. Lastly, delaying *Dnmt1* rescue until after differentiation initiation led to worse outcomes, underscoring the importance of early intervention. Our findings provide foundational insights into the multifaceted mechanisms of DNMT1, emphasize the wide-ranging repercussions of epigenetic perturbations and highlight the challenges associated with rescue interventions.

## MATERIALS AND METHODS

### Cell lines and culture protocol

R1 and *Dnmt1*^tet/tet^ mESCs were gifted by Gregor Andelfinger (University of Montreal) and J. Richard Chaillet (University of Pittsburgh), respectively. Both cell lines were maintained on gelatin-coated plates, free of mouse embryonic fibroblast feeder cells, in mESC culture medium in humidified atmosphere with 5% CO_2_ at 37 °C. Cell culture plates were coated with 0.1% solution of gelatin (Sigma-Aldrich, G1890) in DPBS 1X (Corning, 21-031-CV), then placed in the incubator for 30 minutes after which excess gelatin was removed. Culture medium for mESCs consisted of DMEM (Corning, 10-013-CV) supplemented with 15% ES cell qualified fetal bovine serum (Thermo Fisher, 16141079), 1% 2-mercaptoethanol 100X (Sigma-Aldrich, ES-007-E), 1% non-essential amino acids 100X (Thermo Fisher, 11140050), 1% L-Glutamine 200 mM (Corning, 25-005-CI), 1% nucleosides 100X (Sigma-Aldrich, ES-008-D), 1% penicillin-streptomycin 100X (Corning, 30-002-CI) and 10^3^ units/mL of recombinant mouse LIF protein (Sigma-Aldrich, ESG1107), which was stored at 4 °C for up to 1 week. Culture medium was changed every day and cell confluency was maintained below 80%. Trypsin-EDTA 0.25% (Thermo Fisher, 25200056) was used for passaging cell cultures, typically every other day.

### *Dnmt1* inactivation and rescue in *Dnmt1*^tet/tet^ mESCs

To inactivate *Dnmt1*, *Dnmt1*^tet/tet^ mESCs were treated with 2 μg/mL doxycycline hyclate (Sigma-Aldrich, D9891), a tetracycline derivative, which was added to the culture medium for 6 days. Then, doxycycline treatment was stopped, and cells were maintained for 21 days. Samples were collected prior to doxycycline treatment (*Dnmt1*–control), immediately following doxycycline treatment (*Dnmt1*–inactive), and after the 21-day period of recovery from doxycycline treatment (*Dnmt1*–rescue).

### Protein extraction

Cell pellets were resuspended in lysis buffer, consisting of RIPA buffer with 1mM phenylmethylsulfonyl fluoride (Bio Basic, PB0425), 0.25% Nonidet P-40 (BioShop, NON505), 1X Protease Inhibitor Cocktail (Roche, 11836170001), 20 mM EDTA pH 8.0 (Bio Basic, EB0185), 1X PhosSTOP (Roche, 04906837001). The tubes were then gently shaken for 10 minutes at 4°C and centrifuged at 10000g for 5 minutes at 4°C, after which the supernatants were collected. Proteins were quantified using the PierceTM BCA Protein Assay Kit (Thermo Fisher Scientific, 23227).

### Western blot

Proteins were mixed with Laemmli buffer 5X and loaded into an acrylamide gel (stacking: 4%; resolving: 7%). Migration was performed for 10 minutes at 80 mV followed by 90 minutes at 120 mV until the 30 kDa molecular weight marker (Bio-Rad, 1610374) reached the bottom of the gel. Proteins were then transferred on a nitrocellulose membrane which was saturated under soft agitation for 90 minutes at room temperature with 5% milk (BioShop, SKI400). The membrane was then incubated under soft agitation overnight at 4°C with the primary antibodies for DNMT1 (Cell Signaling Technology, 5032) and β-actin (Cell Signaling Technology, 4970) diluted at 1/1000 in 5% milk. The membrane was washed three times for 10 minutes with TBS-0.1% Tween 20 and incubated under soft agitation for 1 hour at room temperature with the secondary antibody (anti-rabbit IgG HRP-linked; Cell Signaling Technology, 7074) diluted at 1/10000 in 5% milk. The membrane was washed again three times for 10 minutes with TBS-0.1% Tween 20 and revealed using the Clarity Western ECL substrate (Bio-Rad, 1705061) and the ChemiDoc Imaging System.

### Genomic region annotations

Genomic coordinates for known imprinted regions were curated from literature(82–105), sources specific to each region are listed in Table S3. Allele-specific methylated regions in mESCs identified by the Smith Lab(106) in data from Leung et al.(107) were obtained via the UCSC table browser(108) from the DNA methylation Track Hub. Rsubread(109) v2.16.1 was used to obtain gene transcription start sites (TSS), TSS regions (+/- 200 bp from TSS), promoters (−2000 and +200 bp from TSS), and gene bodies. Genomic regions not within promoters or gene bodies were considered as intergenic. Annotatr(110) v1.28.0 was used to obtain CpG islands, CpG island shores (+/- 2kb from CpG islands) and CpG island shelves (+/- 2kb from CpG island shores). Genomic regions not within CpG islands, CpG island shores or CpG island shelves were considered as open sea. Using the UCSC table browser(108), CTCF-bound candidate cis-regulatory elements (cCREs) were extracted from the ENCODE Registry of cCREs(111) filtering for CTCF-only_CTCF-bound, PLS_CTCF-bound, dELS_CTCF-bound, pELS_CTCF-bound and DNase-H3K4me3_CTCF-bound ENCODE classifications. LINEs, SINEs and LTRs were extracted from RepeatMasker using the UCSC table browser(108). The mouse reference genome assembly GRCm38/mm10 was used for all annotations.

### Enzymatic methyl sequencing and data analysis

The following steps were performed in triplicates for each experimental condition: genomic DNA extraction using the QIAamp DNA Mini Kit (QIAGEN, 51304), library preparation using the NEBNext® Enzymatic Methyl-seq Kit (New England Biolabs, E7120L) and 150 bp paired-end sequencing using the Illumina NovaSeq 6000 technology. Library preparation and sequencing was conducted by *Génome Québec*. Between 291M and 385M raw reads were obtained per sample.

Raw reads were processed using the GenPipes(112) methylation sequencing pipeline v3.6.0. Trimmomatic(113) v0.36 was employed for quality trimming of raw reads and removal of sequencing adapters, Bismark(114) v0.18.1 for read alignment to the mouse reference genome assembly GRCm38/mm10, Sambamba(115) v0.7.0 for merging read set files of each sample, Picard v2.9.0 for removing duplicate reads and the methylation extractor function from Bismark(114) v0.18.1 for obtaining the methylation level of single CpGs. Only CpGs that were sequenced in all samples with a coverage ≥ 10X were retained. Methylation levels of single CpGs were then averaged across triplicates.

CpGs were annotated to genomic regions of interest using annotatr(110) v1.28.0. Annotation to promoters was given priority over gene bodies and CpG islands were prioritized over CpG island shores and shelves, with shores taking precedence over shelves. For annotation to transposable elements, only LINEs, SINEs and LTRs with a Smith Waterman alignment score ≥ 1000 were considered. When conducting differential analysis among experimental conditions, we used a 20% change in methylation levels of single CpGs and a 10% change in mean methylation levels across a specific genomic region as thresholds for establishing significance. For comparisons involving groups of multiple CpGs or genomic regions, appropriate statistical testing was applied as described in the quantification and statistical analysis section.

### Chromatin immunoprecipitation sequencing and data analysis

Chromatin immunoprecipitation (ChIP) sequencing was performed in duplicates for each histone modification per experimental condition and one aliquot of non-immunoprecipitated chromatin (i.e. input) was sequenced per experimental condition. First, cells were fixed with a solution consisting of 13% formaldehyde (Sigma-Aldrich, 47608) in DPBS (Corning, 21-031-CV) for 10 minutes at room temperature with gentle rocking. Then, cell and nuclei lysis was performed, and chromatin was purified and sonicated using buffers and protocols from the IDeal ChIP-Seq Histone Kit (Diagenode, C01010051). Sonication was conducted with the Bioruptor® Pico sonication device (Diagenode, B01060010) for 15 cycles (30 seconds ON/ 30 seconds OFF) to obtain DNA fragments between 150-400bp. Next, chromatin was pre-cleared via incubation with protein G Dynabeads™ (Thermo Fisher, 10009D) for 1h at 4°C on a rotator, after which samples were placed on a magnetic rack and input aliquots were taken from the supernatant. Chromatin immunoprecipitation was then conducted as follows. Pre-cleared chromatin samples were incubated overnight at 4°C on a rotator with histone modification antibodies–either anti-H3K4me3 (Active Motif, 39159), anti-H3K27ac (Abcam, ab4729), anti-H3K4me1 (Cell signaling technology, 5326), anti-H3K27me3 (Sigma-Aldrich, 07-449) or anti-H3K9me3 (Abcam, ab8898)–in quantities recommended by the antibody manufacturer. Dynabeads™ were added to the chromatin-antibody mixtures and were incubated for an additional 2h at 4°C on a rotator. Protein A Dynabeads™ (Thermo Fisher, 10001D) or protein G Dynabeads™ (Thermo Fisher, 10009D) were used depending on the antibody source. Washing, elution, decrosslinking and DNA purification were then conducted using the IDeal ChIP-Seq Histone Kit (Diagenode, C01010051). Finally, libraries were prepared with the KAPA HyperPrep Kit (Roche, KK8502) and 50 bp paired-end sequencing was performed at the McGill Genome Centre using the Illumina HiSeq® 2500 technology. Between 16M and 59M raw reads were obtained per sample.

Raw reads were processed using a homemade pipeline including Trimmomatic(113) v0.32 for quality trimming of raw reads and removal of sequencing adapters, Burrows-Wheeler Aligner(116) v0.6.1 for read alignment to the mouse reference genome assembly GRCm38/mm10, the MergeSamFiles.jar and MarkDuplicates.jar scripts from Picard v1.70 for merging read set files of each sample and removing duplicate reads, respectively, and the View function from SAMtools(117) v0.1.18 for filtering reads as to only retain those with a MAPQ score ≥ 15. Then, peak calling was performed using MACS2(118) v2.0.10 with Python v2.7.8, correcting background noise with input signals. Next, MACS2 outputs and BAM files converted to BED format (via the bamTobed function from BEDTools(119) v2.27) were used in the profile_bins function from MAnorm2_utils v1.0.0. The profile_bins function allowed for genomic regions of MACS2 called peaks to be merged into a common set of histone-modified regions across all samples for each histone modification, computing normalized signal enrichment within each region. Finally, MAnorm2(120) v1.0.0 was employed for conducting differential analysis of enrichment signals between experimental conditions, providing a p-value and log_2_ fold change for each histone-modified region. Annotatr(110) v1.28.0 was used to overlap histone-modified regions with other genomic regions and for measuring histone modification RPKM levels within specific regions of interest. Visualization of RPKM signal enrichment within specific target regions in Figure 6E were generated with ComplexHeatmap(121) v2.16.0 and EnrichedHeatmap(122) v1.30.0.

### Mouse sperm and oocyte CpG methylation

Mouse sperm and oocyte CpG methylation profiles were produced by Wang et al.(123) and analyzed by the Smith Lab(106) using the mouse reference genome assembly GRCm38/mm10. These were obtained via the UCSC table browser(108) from the DNA methylation Track Hub and annotated to genomic regions of interest using annotatr(110) v1.28.0 after which mean sperm- and oocyte-specific levels were computed.

### ZFP57 DNA binding motif enrichment analysis

The ZFP57 DNA binding motif MA1583.1(124,125) was obtained from the JASPAR database(126). Enrichment analysis was conducted using the Simple Enrichment Analysis (SEA) tool from the MEME Suite(127) v5.5.5 with shuffled input sequences used as controls.

### mRNA sequencing and data analysis

mRNA sequencing was conducted in triplicates for all experimental conditions. RNA was extracted using the RNeasy Mini Kit (QIAGEN, 74104), libraries were prepared with the NEBNext® Ultra™ II Directional RNA Library Prep Kit (New England Biolabs, E7765) in conjunction with the NEBNext® Poly(A) mRNA Magnetic Isolation Module (New England Biolabs, E7490) and 100 bp paired-end sequencing was performed with the Illumina NovaSeq 6000 technology. Library preparation and sequencing was conducted by *Génome Québec*. Between 27M and 53M raw reads were obtained per sample.

Raw reads were processed using the GenPipes(112) RNA sequencing pipeline v3.1.4. First, Trimmomatic(113) v0.36 was used for quality trimming of raw reads and removal of sequencing adapters and STAR(128) v2.5.3a was used for aligning reads to the mouse reference genome assembly GRCm38/mm10. Then, several functions from Picard v2.9.0 were employed: MergeSamFiles for merging read alignment files of each sample, SortSam for sorting files by coordinate and MarkDuplicates for identifying read duplicates. Next, the HTSeq_count function from HTSeq(129) v0.7.2 was used with Python v2.7.13 for calculating raw counts and StringTie(130) v1.3.5 was used for transcript assembly. Finally, DESeq2(131) v1.18.1 was used with R v3.4.3 for generating normalized counts and conducting differential gene expression analysis between experimental conditions, producing an adjusted p-value and log_2_ fold change for each gene.

### RNA sequencing and data analysis for transposable elements

RNA sequencing was conducted in duplicates for all experimental conditions. The AllPrep DNA/RNA Mini Kit (QIAGEN, 80204) was used for RNA extraction and the TruSeq® Stranded Total RNA Library Prep Human/Mouse/Rat Kit (Illumina, 20020596) for library preparation. 75 bp paired-end sequencing was performed at the McGill Genome Centre with the Illumina HiSeq® 4000 technology. Between 63M and 88M raw reads were obtained per sample.

Raw reads were processed using the GenPipes(112) RNA sequencing pipeline v3.1.4. First, Trimmomatic(113) v0.36 was used for quality trimming of raw reads and removal of sequencing adapters and then reads were aligned to the mouse reference genome assembly GRCm38/mm10 using STAR(128) v2.5.3a. Next, several functions from Picard v2.9.0 were employed including MergeSamFiles for merging read alignment files of each sample, SortSam for sorting files by coordinate and MarkDuplicates for identifying read duplicates. Then, we incorporated the featureCounts(132) function from Subread(133) v2.0.6 to obtain raw counts corresponding to genomic regions of LINE, SINE, and LTR elements, extracted from RepeatMasker using the UCSC table browser(108). Finally, we employed DESeq2(131) v1.18.1 with R v3.4.3 to generate normalized counts for each element, which were then averaged across replicates.

LINEs, SINEs and LTRs were then filtered based on their Smith Waterman alignment score, only retaining those with a score ≥ 1000 for downstream transcriptional analysis. When comparing total normalized counts of an element between experimental conditions, a log_2_ fold change ≤ −1 or ≥ 1 was used as a threshold for establishing significance. For comparisons involving grouped locus-specific normalized counts, appropriate statistical testing was applied as described in the quantification and statistical analysis section.

### Gene set enrichment analysis

Gene set enrichment analysis (GSEA) was conducted using the enricher function from clusterProfiler(134) v4.8.2. All gene sets available for Mus musculus in the Molecular Signatures Database(135,136) were extracted via msigdbr v7.5.1. The following parameters were applied for the GSEA: Benjamini–Hochberg p-value adjustment, minimum gene set size of 10 and maximum gene set size of 1000.

### Visualization of genomic signal tracks

Genomic signal track figures were generated with Gviz(137) v1.44.2. Signal tracks of sample replicates were merged by computing their average signals within common 10-bp bins using the banCoverage function from deepTools(138) v3.5.2, with forward and reverse tracks for RNA-seq and mRNA-seq being priorly merged for each sample. Signal tracks for *Zfp57* WT and KO mESCs were produced by Shi et al.(139) (GSE123942) and for *Setdb1* WT and KO mESCs by Barral et al.(140)(GSE171749), which were obtained via the Gene Expression Omnibus(141,142).

### FACS assay for detecting γH2AX and assessing cell size and granularity

The H2AX phosphorylation (γH2AX) for fluorescence-activated cell sorting (FACS) kit (Sigma-Aldrich, 17-344) was used to prepare cell samples following the manufacturer’s protocol for fixation, permeabilization and staining. All experimental conditions were assayed in triplicates for γH2AX, and one unstained control sample was included per condition. One DNA damage positive control sample stained for γH2AX was also included, which consisted of *Dnmt1**^CTL^*** mESCs that were irradiated with 10 Gγ using a Faxitron® irradiator six days prior to sorting. Samples were sorted with the BD LSRFortessa™ Cell Analyzer (BD Biosciences) and data was processed with the BD FACSDiva™ software v8.0.2. Forward scatter area (FSC-A) and side-scatter area (SSC-A) measurements were used for filtering out apoptotic-like cells prior to fluorescence analysis. FSC-A and SSC-A measurements of the live cell population were then used for comparing cell size and granularity, respectively, across experimental conditions.

### Measuring relative telomere length by qPCR

Genomic DNA was extracted from *Dnmt1^t^*^et/tet^ mESCs and R1 mESCs using the QIAamp DNA Mini Kit (QIAGEN, 51304). qPCR primer sequences for mouse telomeric and *Rplp0* (single-copy gene) DNA were obtained from Callicott & Womack(143), who adapted the protocol developed by Cawthon(144) for measuring relative telomere length in human samples. qPCR primers were ordered from Integrated DNA Technologies and evaluated using customary PCR techniques, including annealing temperature optimization and amplification efficiency assessment. qPCR assays were performed using a LightCycler® 480 instrument (Roche) and the SensiFAST™ SYBR® No-ROX Kit (Bioline, BIO- 98005), following the kit’s 3-step protocol for 30 cycles with an annealing temperature of 62 °C. Telomeric and *Rplp0* DNA were amplified by qPCR in all *Dnmt1^t^*^et/tet^ mESC conditions and in R1 mESCs, with eight replicates for each amplification to ensure reproducibility. For telomeric DNA amplifications, 0.2 ng of DNA was used and for *Rplp0* DNA amplifications, 2 ng of DNA was used. Relative telomere length was calculated for individual replicates of each *Dnmt1^t^*^et/tet^ mESC condition with the delta-delta Ct method(145) as in the protocol by Cawthon, using *Rplp0* as the reference single-copy gene for normalization and R1 mESCs as the reference DNA sample for relativization. The average *Rplp0* Ct value for each condition was used for normalizing telomeric DNA Ct values and the average normalized telomeric DNA Ct value in R1 mESCs was used to relativize normalized telomeric DNA Ct values in *Dnmt1^t^*^et/tet^ mESCs.

- Mouse telomeric DNA forward primer sequence: CGGTTTGTTTGGGTTTGGGTTTGGGTTTGGGTTTGGGTT
- Mouse telomeric DNA reverse primer sequence: GGCTTGCCTTACCCTTACCCTTACCCTTACCCTTACCCT
- Mouse *Rplp0* DNA forward primer sequence: ACTGGTCTAGGACCCGAGAAG
- Mouse *Rplp0* DNA reverse primer sequence: TCAATGGTGCCTCTGGAGATT

### Assessment of cell proliferation rate

The Incucyte® S3 Live-Cell Imaging and Analysis System and its associated software for data processing (Sartorius) were used to assess cell proliferation rate by measuring the increase in confluency over the course of 30 hours. Cells were seeded in 6-well plates at a density of 50,000 cells/cm^2^ in triplicates for all experimental conditions and confluency measurements were taken at 0, 6, 12, 18, 24 and 30 hours.

### Differentiation of mESCs towards endoderm, mesoderm, and ectoderm lineages

After thawing mESCs, they were passaged twice according to the mESC culture protocol. The day before beginning differentiation protocols, mESCs were passaged using TrypLE™ Express Enzyme 1X (Thermo Fisher, 12604021) supplemented with 10 μM ROCK Inhibitor Y-27632 (STEMCELL Technologies, 72302) for detachment, re-seeding them with a density of 30,000 cells/cm^2^ in mESC culture medium in dishes coated with laminin from mouse Engelbreth-Holm-Swarm sarcoma (Roche, 11243217001). For laminin coating, 2.5 μg/cm^2^ of laminin diluted in DMEM (Corning, 10-013-CV) was added to cell culture dishes in a minimal volume to coat entire dish surface, after which excess laminin was removed, and dishes were air dried for 45 minutes inside sterile cell culture hood. The next day, mESC cultures were rinsed twice with differentiation medium prior to starting differentiation protocols. Differentiations were conducted in triplicates for each experimental condition, maintaining cultures in humidified atmosphere with 5% CO_2_ at 37 °C without passaging. Differentiation cultures were rinsed twice daily with differentiation medium prior to adding differentiation treatments to remove dead cells as a fair amount of cell death was expected. Differentiation treatment mixtures were freshly prepared each day.

For endoderm differentiation, the differentiation medium consisted of RPMI (Thermo Fisher, 11875093) supplemented with 10% KnockOut™ Serum Replacement (Thermo Fisher, 10828028), 2% B-27™ minus insulin 50X (Thermo Fisher, A1895601), 1.5% HEPES 1M (Thermo Fisher, 15630106), 1% 2-mercaptoethanol 100X (Sigma-Aldrich, ES-007-E), 1% non-essential amino acids 100X (Thermo Fisher, 11140050), 1% L-Glutamine 200 mM (Corning, 25-005-CI), 1% nucleosides 100X (Sigma-Aldrich, ES-008-D) and 1% penicillin-streptomycin 100X (Corning, 30-002-CI). Cells were treated with 5 μM CHIR99021 (Sigma-Aldrich, SML1046) and 50 ng/mL Activin A (R&D Systems, 338-AC) for 1 day followed by 50 ng/mL Activin A and 100 nM LDN193189 (STEMCELL Technologies, 72147) for 2 days.

For mesoderm differentiation, the differentiation medium consisted of EMEM (Corning, 10009CV), 10% KnockOut™ Serum Replacement (Thermo Fisher, 10828028), 1% sodium pyruvate 100 mM (Thermo Fisher, 11360070), 1% 2- mercaptoethanol 100X (Sigma-Aldrich, ES-007-E), 1% non-essential amino acids 100X (Thermo Fisher, 11140050), 1% L-Glutamine 200 mM (Corning, 25-005-CI), 1% nucleosides 100X (Sigma-Aldrich, ES-008-D) and 1% penicillin-streptomycin 100X (Corning, 30-002-CI). Cells were treated with 5 μM CHIR99021 (Sigma-Aldrich, SML1046) and 21 ng/mL BMP-4 (R&D Systems, 5020-BP) for 3 days.

For ectoderm differentiation, the differentiation medium was the same as the mESC culture medium but without recombinant mouse LIF protein. Cells were treated with 1 μM of retinoic acid (Sigma-Aldrich, R2625) for 3 days.

On the last day of differentiation protocols, differentiation cultures were sorted based on lineage-specific cell surface and viability markers using fluorescence-activated cell sorting (FACS). Differentiation cultures were rinsed twice with FACS buffer consisting of DPBS 1X (Corning, 21-031-CV) supplemented with 1% ES cell qualified fetal bovine serum (Thermo Fisher, 16141079). A minimal volume of FACS buffer was added to coat entire dish surface and cells were dissociated into single-cell suspension with a cell scraper and gentle pipetting and were transferred to conical tubes. Primary antibodies for cell surface markers were added to cell suspensions in the amount recommended by the manufacturer followed by an incubation of 45 minutes at 4°C, gently mixing every 10 mins. The primary antibodies used for endoderm, mesoderm and ectoderm were anti-CXCR4 (R&D Systems, MAB21651), anti-CDH2 (R&D Systems, AF6426) and anti-NCAM1 (R&D Systems, AF2408), respectively. Cell suspensions were then rinsed twice with FACS buffer and resuspended in FACS buffer containing the fluorophore-conjugated secondary antibody in the amount recommended by the manufacturer followed by an incubation of 45 minutes at 4°C, gently mixing every 10 mins. The secondary antibodies used for endoderm, mesoderm and ectoderm were APC-conjugated anti-Rat IgG (R&D Systems, F0113), APC-conjugated anti-Sheep IgG (R&D Systems, F0127) and Alexa Fluor™ 488-conjugated anti-Goat IgG (Thermo Fisher, A11055), respectively. Cell suspensions were rinsed twice with FACS buffer and resuspended in FACS buffer at a density of 2M cells/100 μL, transferring them to FACS tubes and adding 5 μL per sample of propidium iodide 10 μg/mL (Thermo Fisher, P1304MP), a viability marker, which was used for retaining only live cells. Cells were sorted with the BD FACSAria™ Fusion Flow Cytometer (BD Biosciences) and data was processed with the BD FACSDiva™ software v8.0.2. One unstained control sample per differentiation condition and one control sample of dead cells stained with propidium iodide were also sorted. The control sample of dead cells consisted of pooled differentiated cells heated at 65°C for 1 minute in a digital dry bath and then incubated on ice for 1 minute. After sorting, RNA was immediately extracted and stored at −80°C for subsequent mRNA sequencing.

### Embryoid body formation

*Dnmt1*^tet/tet^ mESCs (*Dnmt1*–control, *Dnmt1*–inactive and *Dnmt1*–rescue conditions) were detached with TrypLE™ Express Enzyme 1X (Thermo Fisher, 12604021) supplemented with 10 μM ROCK Inhibitor Y-27632 (STEMCELL Technologies, 72302) and counted with a hemocytometer. mESCs were seeded into 96-well round bottom ultra-low attachment plates (Corning, 7007) at 1000 cells per well in 150 μL of culture medium, centrifuged at 100g for 3 minutes and left undisturbed for 2 days. The culture medium used for the first 2 days of embryoid body formation was the same as the mESC culture medium but without nucleosides and with 10 μM ROCK Inhibitor Y-27632. From day 2 to day 4, the embryoid body culture medium was the same as the mESC medium but without LIF and nucleosides. From day 4 to day 10, the embryoid body culture medium was the same as the mESC medium but without LIF and nucleosides and with DMEM/F12 (Thermo Fisher, 11330032) instead of DMEM. Culture medium was changed daily by replacing two thirds with fresh medium. Doxycycline hyclate (2 μg/mL; Sigma-Aldrich, D9891) was added to the embryoid body culture medium from day 0 (seeding) to day 10 for the *Dnmt1*–inactive condition, or from day 0 to day 5 for the post-differentiation *Dnmt1*–rescue condition. For post-differentiation rescue of *Dnmt1*, embryoid bodies were rinsed three times on day 5 with culture medium to remove doxycycline. Embryoid body formation thus included four experimental conditions: *Dnmt1*–control, *Dnmt1*–inactive, *Dnmt1*–rescue and post-differentiation *Dnmt1*–rescue. Embryoid bodies (n=192 per condition) were derived from the same *Dnmt1*^tet/tet^ mESC culture. From day 2 to day 10 of embryoid body formation, daily bright-field microscopy images were taken of 20 embryoid bodies per condition with an EVOS M5000 Cell Imaging System (Thermo Fisher). These images were used for measuring embryoid body surface area and perimeter with ImageJ software version 1.54g. Surface area was used as an indicator of size and circularity was determined by the formula 4π × Area/Perimeter^2^. Embryoid bodies were harvested and pooled on days 5 and 10 for protein and RNA extractions for Western blotting and mRNA sequencing.

### Deconvolution of embryoid body mRNA sequencing data

Raw mRNA-seq counts of embryoid bodies were deconvoluted from single-cell RNA-seq data of mouse gastrulation embryos obtained from the R package MouseGastrulationData v1.18.0(146), using only data from embryonic day 7.75 embryos and selecting only extraembryonic (ExE) endoderm, epiblast, primitive streak, rostral neurectoderm and forebrain/midbrain/hindbrain cell types. Rostral neurectoderm and forebrain/midbrain/hindbrain cell types were combined into one cell type category that we named neuroectoderm. Technical artifacts were filtered out of the single-cell data by removing doublets and cells with stripped nuclei. The deconvolution was performed with the function music2_prop_t_statistics from the R package MuSiC v1.0.0(147,148) using default function settings.

### Quantification and statistical analysis

Statistical tests for comparing grouped data were conducted in R v4.3.1 using functions from the stats R core package. To determine whether parametric or non-parametric statistical tests should be used, we first applied the Shapiro-Wilk test for normality using the shapiro.test function as well as Levene’s test for homogeneity of variance using the leveneTest function. Parametric testing was conducted if both the Shapiro-Wilk and Levene’s tests passed, whereas non-parametric testing was conducted if either one failed. For pairwise comparisons across more than two experimental conditions, parametric testing was performed using one-way ANOVA followed by Tukey’s HSD test with p-value adjustment via the aov and TukeyHSD functions and non-parametric testing was performed using the Kruskal-Wallis test followed by the Wilcoxon rank-sum test with Benjamini-Hochberg p-value adjustment via the kruskal.test and pairwise.wilcox.test functions. For comparing two groups of data, parametric testing was performed with Student’s t test using the t.test function, and non-parametric testing was performed with the Kruskal-Wallis test using the kruskal.test function. Pearson’s chi-squared tests were performed with the chisq.test function and principal component analysis with the prcomp function. Differential quantification and statistical analyses of histone-modified regions using MAnorm2 and of gene expression using DESeq2 are described in their respective method details section.

## RESULTS

### Inactivation and rescue of *Dnmt1* in mESCs induces extensive epigenomic alterations

Utilizing *Dnmt1*^tet/tet^ mESCs(36), we previously revealed through reduced-representation bisulfite sequencing (RRBS) that loss of DNA methylation (i.e. CpG methylation) due to *Dnmt1* inactivation is not fully restored upon *Dnmt1* rescue(37). However, RRBS can only capture ∼10% of the total CpGs in the genome, with considerable bias toward CpG-dense promoter regions(149–153). To achieve a comprehensive genome-wide view of DNA methylation alterations throughout *Dnmt1* inactivation and rescue, here we employed the *Dnmt1*^tet/tet^ mESC model and performed enzymatic methyl sequencing (EM-seq) which provides vastly superior CpG coverage and methylation sensitivity. In this study, 15,349,152 CpGs (∼73% of total CpGs) were sequenced with EM-seq compared to 1,422,634 CpGs (∼7% of total CpGs) in our previous study with RRBS. We also conducted genome-wide chromatin immunoprecipitation sequencing (ChIP-seq) for five key histone modifications (permissive: H3K4me3, H3K27ac and H3K4me1; repressive: H3K27me3 and H3K9me3) to investigate the broader impact on epigenomic landscapes. Herein, untreated *Dnmt1*^tet/tet^ mESCs are referred to as *Dnmt1*–control (*Dnmt1**^CTL^***), those after *Dnmt1* inactivation by doxycycline as *Dnmt1*–inactive (*Dnmt1**^INV^***), and those after rescuing *Dnmt1* through doxycycline removal as *Dnmt1*–rescue (*Dnmt1**^RES^***) (Figure 1A). Successful inactivation and rescue of *Dnmt1* were confirmed by Western blot (Figure S8A).

**Figure 1:**
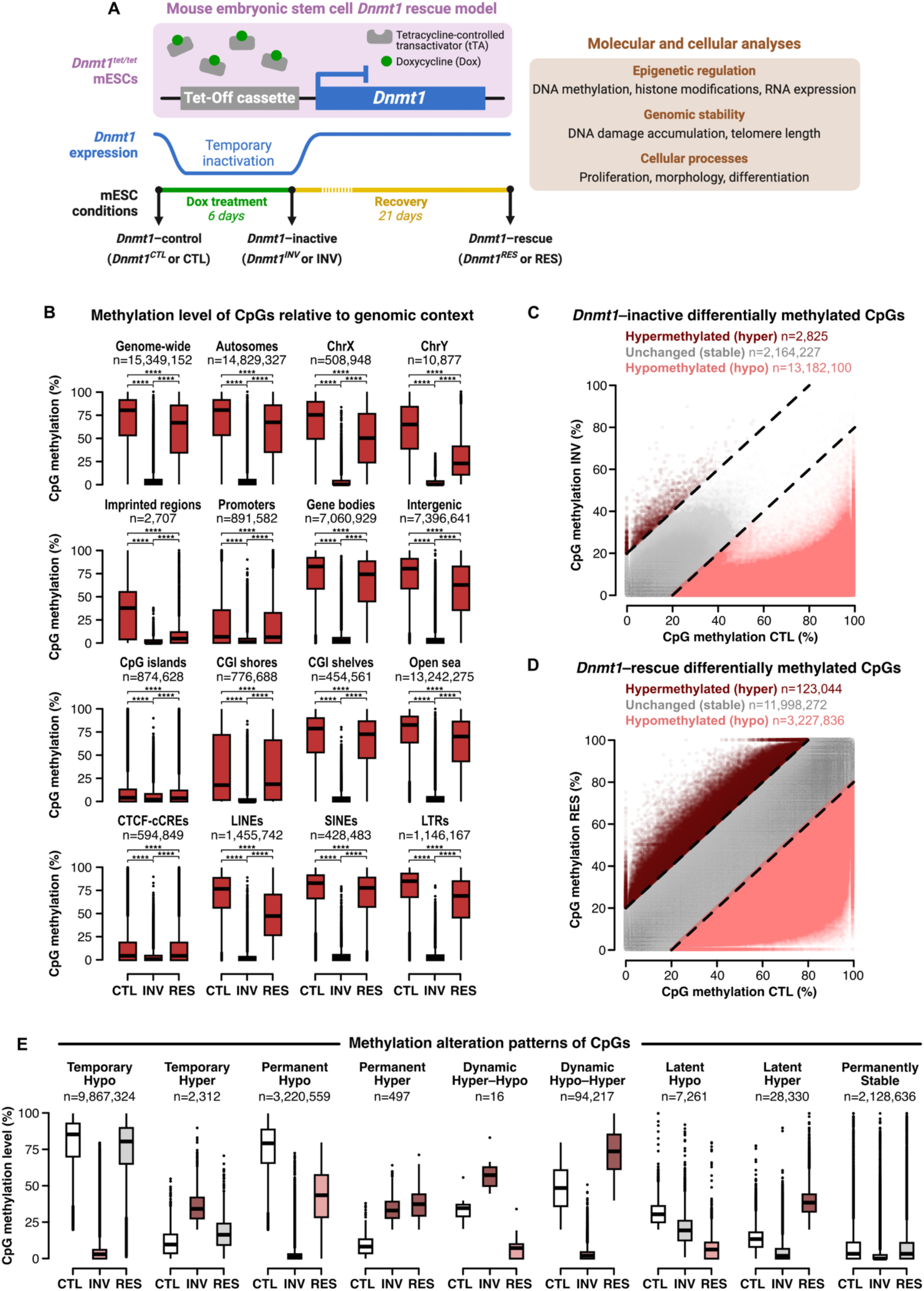
*Dnmt1* inactivation and rescue permanently reduces CpG methylation levels across genomic contexts, with over 3.2 million permanently hypomethylated single CpGs. **(A)** Schematic diagram depicting the *Dnmt1*^tet/tet^ mESC model and molecular and cellular analyses performed. Made with Biorender.com. **(B)** Pairwise comparisons of single CpG methylation levels in *Dnmt1^CTL^*, *Dnmt1^INV^* and *Dnmt1^RES^* (EM-seq: n=3 per condition) relative to genomic location using the Wilcoxon rank sum test with Benjamini-Hochberg p-value adjustment. **(C-D)** Differential methylation level of genome-wide single CpGs in *Dnmt1^INV^* and *Dnmt1^RES^*versus *Dnmt1^CTL^*. Hypermethylated: increase ≥ 20%. Hypomethylated: decrease ≥ 20%. **(E)** Methylation alteration patterns of single CpGs during *Dnmt1* inactivation and rescue determined by differential states in C and D. Temporary: hypo/hyper in *Dnmt1^INV^*& stable in *Dnmt1^RES^*. Permanent: hypo/hyper in both *Dnmt1^INV^*& *Dnmt1^RES^*. Dynamic: hypo/hyper in *Dnmt1^INV^* & the opposite in *Dnmt1^RES^*. Latent: stable in *Dnmt1^INV^* & hypo/hyper in *Dnmt1^RES^*. Permanently stable: stable in both *Dnmt1^INV^* & *Dnmt1^RES^*.

We first compared overall methylation levels of single CpGs in *Dnmt1**^CTL^**, Dnmt1**^INV^***, and *Dnmt1**^RES^*** conditions across various genomic location contexts, revealing a near-total depletion in *Dnmt1**^INV^*** and a sustained decrease in *Dnmt1**^RES^*** for all contexts (Figure 1B). Permanent reduction in CpG methylation levels observed in *Dnmt1**^RES^***was particularly sizable for chromosome X (−17%), chromosome Y (−32%), imprinted regions (−27%), intergenic (−13%), open sea (− 10%), LINEs (−21%), and LTRs (−13%), in contrast to other genomic location contexts (genome-wide: −9%, autosomes: −9%, promoters: −2%, gene bodies: −6%, CpG islands: −0.6%, CGI shores: −1%, CGI shelves: −4%, CTCF-bound candidate cis-regulatory elements (cCREs): −0.4%, SINEs: −5%). However, considering that CpGs in promoters, CpG islands, and CTCF-bound cCREs were mostly poorly methylated in *Dnmt1**^CTL^*** (median level < 7%), the extent of their permanent hypomethylation may not be directly comparable to regions with initially highly methylated CpGs.

To explore genome-wide methylation dynamics of single CpGs throughout *Dnmt1* inactivation and rescue, we measured the change in methylation for each CpG in *Dnmt1**^INV^*** versus *Dnmt1**^CTL^*** (Figure 1C) and in *Dnmt1**^RES^*** versus *Dnmt1**^CTL^*** (Figure 1D). We characterized a methylation increase ≥ 20% as hypermethylated and a decrease ≥ 20% as hypomethylated; all other changes were considered as stably methylated. Each CpG was then categorized based on its methylation change (hypermethylated, hypomethylated, stable) in *Dnmt1**^INV^*** and *Dnmt1**^RES^***, revealing nine alteration patterns (Figure 1E). Among these, temporary hypomethylation, permanent hypomethylation, and permanent stability of methylation were the most prevalent, comprising ∼9.9 million, ∼3.2 million, and ∼2.1 million CpGs, respectively. Although permanent stability was prominent, those CpGs were mostly poorly methylated in *Dnmt1**^CTL^*** (median level < 3.4%), indicating limited potential for hypomethylation, unlike temporarily or permanently hypomethylated CpGs which were mostly highly methylated in *Dnmt1**^CTL^*** (median level > 85% and 79%, respectively). Some CpG hypermethylation was observed, primarily in a dynamic hypo–hyper pattern (n=94,217) or a latent hypermethylation pattern (n=28,330). The remaining five patterns (temporary hyper, permanent hyper, dynamic hyper–hypo, latent hypo) contained a negligible number of CpGs, totaling ∼10K.

Next, we investigated the impact of *Dnmt1* inactivation and rescue in mESCs on histone modification landscapes. We first performed peak calling for H3K4me3, H3K27ac, H3K4me1, H3K27me3 and H3K9me3 in each condition. Compared to *Dnmt1**^CTL^***, the total number of peaks was higher in *Dnmt1**^INV^*** for all modifications, while lower peak numbers were observed in *Dnmt1**^RES^***, except for H3K4me3 which remained higher (Figure S1A). Recognizing that peak numbers alone may not fully capture fluctuations in histone modification levels due to variable peak size and enrichment, peak genomic regions across experimental conditions were merged into a common set of regions for each histone modification while also conducting signal enrichment normalization. This enabled statistical and quantitative comparisons between experimental conditions for detecting differentially histone-modified regions (DHMs). DHMs exhibiting either a decrease or an increase in histone modification enrichment, and regions that maintained stable enrichment were computed in *Dnmt1**^INV^*** (Figure 2A) and in *Dnmt1**^RES^*** (Figure 2B) versus *Dnmt1**^CTL^***. Subsequently, histone-modified regions were classified into histone modification alteration patterns based on their differential states (decrease, increase or stable) in *Dnmt1**^INV^*** and *Dnmt1**^RES^*** conditions (Figures 2C and S1B). Although most regions displayed permanent stability of histone modification enrichment (Figure S1B), thousands of DHMs were detected in *Dnmt1**^INV^***and *Dnmt1**^RES^*** (Figures 2A-B), with H3K4me3 increase in *Dnmt1**^INV^***being the most striking. Interestingly, DHMs were predominantly condition-specific; the majority of DHMs in *Dnmt1**^INV^*** exhibited temporary alteration (Figure 2D) and in *Dnmt1**^RES^*** showed latent alteration (Figure 2E).

**Figure 2:**
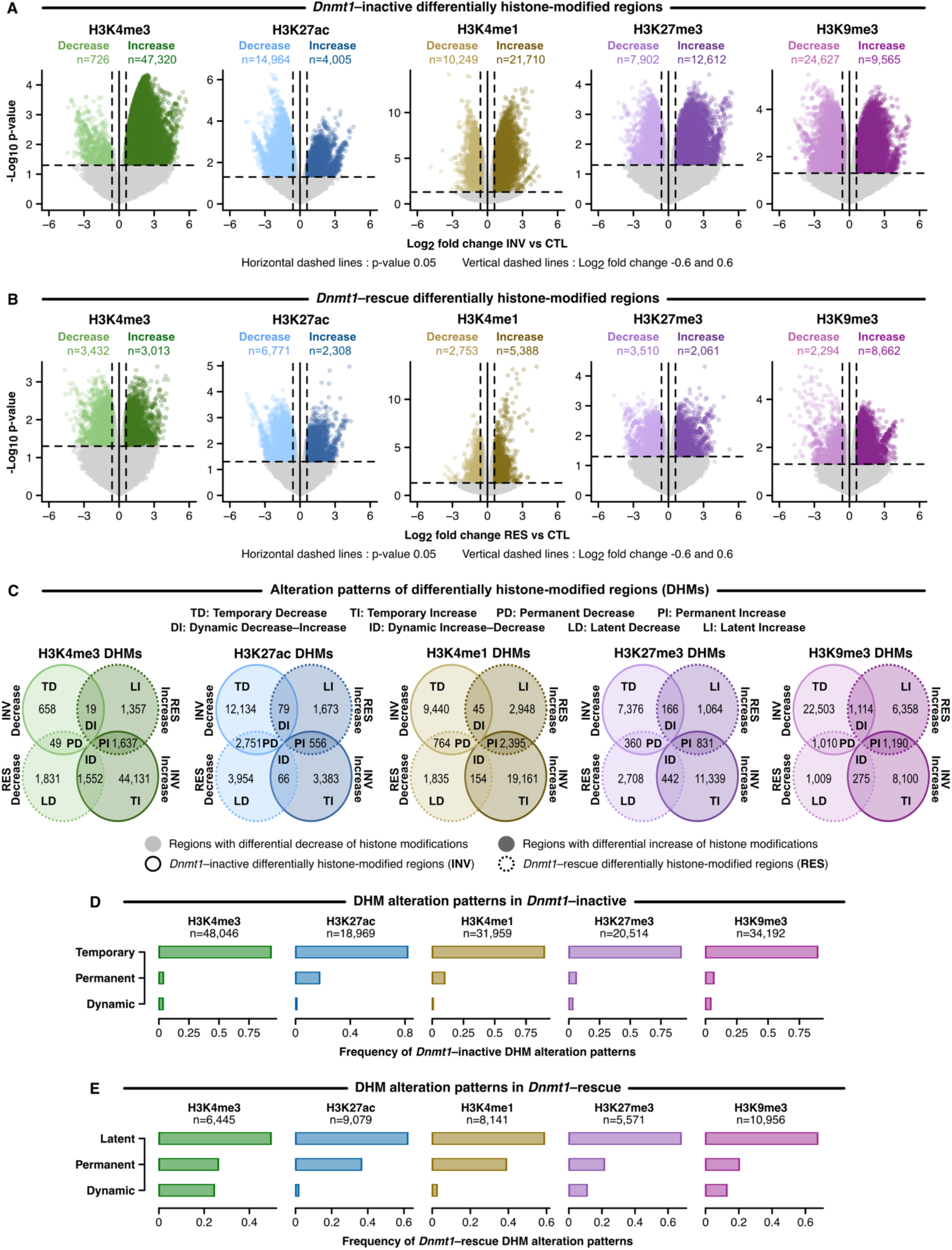
*Dnmt1* inactivation and rescue in mESCs alters genome-wide histone modification landscapes. **(A-B)** Differentially histone-modified regions (DHMs) for H3K4me3, H3K27ac, H3K4me1, H3K27me3 and H3K9me3 in *Dnmt1**^INV^*** and *Dnmt1**^RES^*** versus *Dnmt1**^CTL^*** measured by ChIP sequencing (n=2 per condition) and determined by a p- value ≤ 0.05 and log_2_ fold change ≤ −0.6 or ≥ 0.6 using the MAnorm2 software. **(C)** Alteration patterns of DHMs for each histone modification throughout *Dnmt1* inactivation and rescue determined by their differential state in A and B. Temporary: decrease/increase in *Dnmt1**^INV^*** & stable in *Dnmt1**^RES^***. Permanent: decrease/increase in both *Dnmt1**^INV^*** & *Dnmt1**^RES^***. Dynamic: decrease/increase in *Dnmt1**^INV^*** & the opposite in *Dnmt1**^RES^***. Latent: stable in *Dnmt1**^INV^***& decrease/increase in *Dnmt1**^RES^***. **(D-E)** Frequency of DHM alteration patterns for *Dnmt1**^INV^*** and *Dnmt1**^RES^***, respectively, representing the number of DHMs in each pattern divided by the total number of DHMs. *See also Figure S1*.

In summary, DNA methylation and histone modification landscapes were extensively altered throughout *Dnmt1* inactivation and rescue, showing that the effects of transient loss of DNMT1 activity extends beyond DNA methylation to affect broader epigenomic networks. This reveals the critical role of sustained DNA methylation maintenance for preserving epigenomic integrity across multiple layers, emphasizing the importance of considering wider epigenomic contexts when key epigenetic mechanisms are disrupted. Although most alterations triggered by *Dnmt1* inactivation were reversed upon rescue, many alterations persisted, and additional alterations emerged post-rescue, illustrating the challenges associated with fully rescuing epigenetic perturbations without collateral effects.

### Permanent CpG hypomethylation is associated with specific genomic location contexts and histone modification alteration patterns

*Dnmt1* inactivation and rescue permanently reduced DNA methylation levels of over 3.2 million CpGs. To elucidate underlying factors, we examined whether specific genomic contexts or certain histone modification alteration patterns were more or less associated with permanent hypomethylation of these CpG sites.

First, we computed the number of CpGs with the potential to undergo permanent hypomethylation, i.e. a sustained methylation decrease of at least 20% throughout *Dnmt1**^INV^*** and *Dnmt1**^RES^***. These CpGs had to exhibit a methylation level exceeding 20% in the *Dnmt1**^CTL^***condition and were designated as methylated CpGs (mCpGs). The frequency of mCpGs (ratio of mCpGs to sequenced CpGs) was high (≥ 0.65) across most genomic contexts, except for promoters (0.33) and CGI shores (0.49), which exhibited moderate frequencies, and CpG islands (0.15) and CTCF-bound cCREs (0.24), where frequencies were markedly lower (Figure S2A; Table S1). As for histone modification alteration patterns, H3K4me1 and H3K9me3 patterns exhibited high mCpG frequencies overall, while H3K4me3, H3K27ac and H3K27me3 patterns had variable mCpG frequencies (Figure S2B; Table S2). Next, we quantified the number of permanently hypomethylated CpGs and their frequency (ratio of permanently hypomethylated CpGs to mCpGs) (Figures 3A and 3C; Tables S1 and S2). This was followed by the application of Pearson’s chi-squared test to discern positive, negative, or neutral association of permanent CpG hypomethylation with genomic location contexts (Figure 3B; Table S1) or with histone modification alteration patterns (Figure 3D, Table S2). Positive associations were observed for imprinted regions, sex chromosomes, LINEs, LTRs, intergenic, and open sea genomic location contexts. All H3K9me3 patterns, except dynamic increase–decrease, also exhibited positive association along with permanent and latent increase patterns of H3K4me3 and H3K27me3. While the permanent increase pattern of H3K27ac displayed neutral association, it approached significance for positive association. It is noteworthy that, despite having high mCpG frequencies in *Dnmt1**^CTL^***, certain genomic location contexts (e.g. gene bodies, CGI shelves, SINEs) and histone modification alteration patterns (e.g. latent decrease of H3K4me3, temporary increase of H3K4me1, H3K27ac or H3K27me3) were negatively associated with permanent CpG hypomethylation.

**Figure 3:**
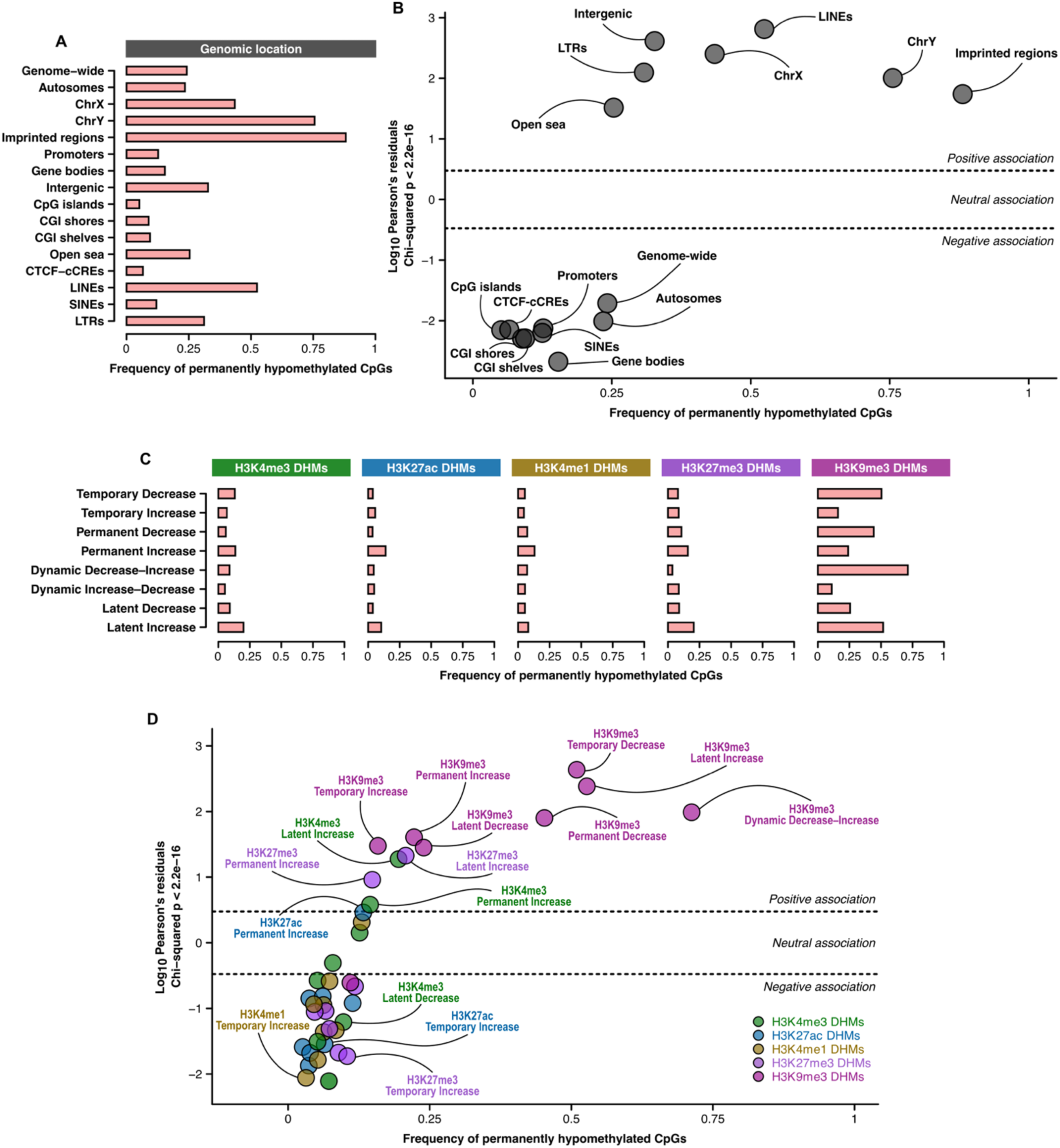
Permanent CpG hypomethylation is associated with specific genomic location contexts and histone modification alteration patterns. **(A)** Frequency of permanently hypomethylated CpGs per genomic location context. **(B)** Pearson’s chi-squared test associating the frequency of permanently hypomethylated CpGs with genomic location contexts. **(C)** Frequency of permanently hypomethylated CpGs in differentially histone-modified regions (DHMs) categorized into alteration patterns identified in Figure 2C. **(D)** Pearson’s chi-squared test associating the frequency of permanently hypomethylated CpGs with DHM alteration patterns. Frequency of permanently hypomethylated CpGs represents the number of permanently hypomethylated CpGs divided by the number of methylated (methylation level ≥ 20%) CpGs in *Dnmt1^CTL^*. Pearson’s residual ≥ 2 indicates positive, < 2 but > −2 indicates neutral and ≤ −2 indicates negative association. *See also Tables S1 and S2 and Figure S2*.

In brief, specific genomic contexts, including imprinted regions, LINEs, LTRs and sex chromosomes were more susceptible to permanent CpG hypomethylation during *Dnmt1* inactivation and rescue. These results reinforce the necessity of sustained DNMT1 activity to ensure the heritability of DNA methylation at imprinted loci(36,37) and suggest that other non-imprinted regions may possess heritable DNA methylation marks, validating findings from our previous study using RRBS(37). Additionally, permanent CpG hypomethylation being positively associated with sex chromosomes could relate to how epigenetic perturbations often result in sex-specific consequences(11,23). Specific patterns of histone modification alterations, particularly H3K9me3, were also positively associated, suggesting that permanent loss of DNA methylation involved additional changes to the chromatin microenvironment. Therefore, simply reinstating DNA methylation maintenance is possibly insufficient to restore DNA methylation levels in some regions; concurrent targeting of histone modifications may be required to fully rescue epigenetic states.

### Analysis of known imprinted regions enables the identification of 20 regions with imprinted-like epigenetic and regulatory signatures

Genomic imprinting is an epigenetic phenomenon whereby certain genes are expressed in a parent-of-origin manner(154). Epigenetic disruptions can cause failure of genomic imprinting maintenance, leading to severe developmental disorders(155,156). Imprinted regions are CpG-dense long-range cis-acting regulatory sites characterized by heritable allele-specific DNA methylation, maintained by the concerted actions of DNMT1, ZFP57, TRIM28 and SETDB1-mediated deposition of broad H3K9me3 domains(35–37,107,139,154,157,158). Although genomic imprinting remains an active field of research, its heritability mechanisms are not fully understood, and the search for novel imprinted regions is still ongoing. Here, we leveraged *Dnmt1*^tet/tet^ mESCs to better understand how histone modifications contribute to genomic imprinting heritability and uncover other regions with imprinted signatures.

Among the 26 imprinted regions(82–105) that were examined, 15 had permanently hypomethylated CpGs, exhibited permanent mean CpG hypomethylation ≥ 10% and overlapped permanently decreased H3K9me3 regions (Figure 2C), specifically within broad H3K9me3 domains (≥ 3000 bp) (Figure 4A; Table S3). The remaining 11 did not meet those criteria in addition to having low mCpG frequencies (≤ 0.25) and low mean CpG methylation levels in *Dnmt1**^CTL^*** (median < 4%), thus were excluded from further analysis. To identify regions with imprinted-like signatures, we employed publicly available mESC allele-specific methylated regions identified by the Smith Lab(106) in data from Leung et al.(107) We filtered allele-specific methylated regions based on their non-overlap with known imprinted regions and on them exhibiting epigenetic dynamics akin to those observed in the 15 known imprinted regions selected in Figure 4A: presence of permanently hypomethylated CpGs, permanent mean CpG hypomethylation ≥ 10% and overlapping permanently decreased H3K9me3 regions within broad domains ≥ 3000 bp. Ultimately, 20 imprinted-like regions were retained, named according to their intersection with or proximity to genes (Figure 4B; Table S3). In recent work by Yang et al.(159), genomic regions with imprinted signatures were identified in mouse preimplantation embryos through allele-specific profiling of DNA methylation and H3K9me3 alongside global depletion of H3K9me3 to establish mechanistic co-dependency. Their analysis unveiled 22 imprinted-like regions and performed Cas9-mediated deletions of five of them, demonstrating their importance for mouse development. Compellingly, three of our imprinted-like regions—*Spred2*, *Lipt1-Mitd1* and *Zfp668*—coincided with imprinted-like regions identified by Yang et al. (Figure S3A), with *Lipt1-Mitd1* among the five that were confirmed biologically relevant.

**Figure 4:**
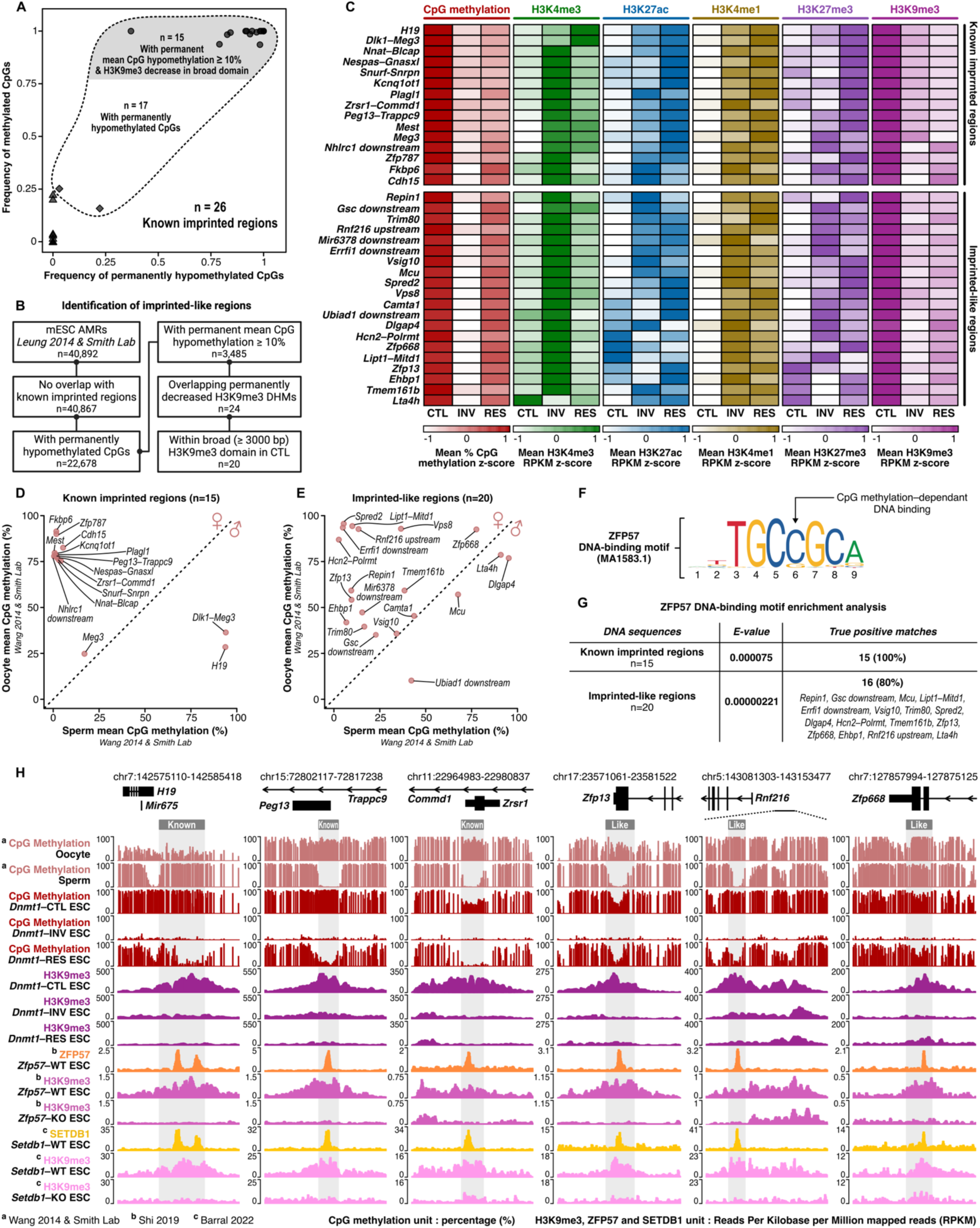
Analysis of known imprinted regions enables the identification of 20 regions with imprinted-like epigenetic and regulatory signatures. **(A)** Selection of 15 known imprinted regions based on four criteria: permanently hypomethylated CpGs ≥ 1, permanent mean CpG hypomethylation ≥ 10%, overlapping permanently decreased H3K9me3 DHMs and located within H3K9me3 broad domains (≥ 3000 bp) in *Dnmt1^CTL^*. **(B)** Workflow for identifying imprinted-like regions among mESC allele-specific methylated regions based on the four criteria in A. **(C)** Z-score normalization of epigenetic modification levels in *Dnmt1^CTL^*, *Dnmt1^INV^*and *Dnmt1^RES^* for selected known imprinted regions and imprinted-like regions. **(D-E)** Mean CpG methylation levels in oocytes versus sperm for selected known imprinted regions and imprinted-like regions, respectively. **(F)** ZFP57 DNA binding motif MA1583.1. **(G)** Enrichment of ZFP57 DNA binding motif MA1583.1 in selected known imprinted regions and imprinted-like regions. **(H)** Genomic signal tracks for *H19*, *Peg13–Trappc9* and *Zrsr1–Commd1* imprinted regions and *Zfp13*, *Rnf216 upstream* and *Zfp668* imprinted-like regions showing CpG methylation in gametes and *Dnmt1*^tet/tet^ mESCs, H3K9me3 in *Dnmt1*^tet/tet^ mESCs, *Zfp57* WT and KO mESCs and *Setdb1* WT and KO mESCs, ZFP57 binding in *Zfp57* WT mESCs and SETDB1 binding in *Setdb1* WT mESCs. *See also Table S3 and Figures S3 and S4*.

In comparison to the selected 15 known imprinted regions, imprinted-like regions also had high frequencies of mCpGs in *Dnmt1**^CTL^*** and high frequencies of permanently hypomethylated CpGs (Figure S3B), along with comparable mean CpG methylation levels in *Dnmt1**^CTL^*** (Figure S3C). However, imprinted-like regions displayed lower CpG density (Figure S3D) and less severe permanent mean CpG hypomethylation (Figure S3E) than the selected 15 known imprinted regions. Next, we computed mean histone modification RPKM levels within each region in *Dnmt1**^CTL^***, *Dnmt1**^INV^***and *Dnmt1**^RES^*** (Table S3), employing z-score normalization to reveal epigenetic alteration dynamics other than permanent CpG hypomethylation and H3K9me3 decrease (Figure 4C). Notably prevalent among known imprinted regions and imprinted-like regions were a temporary sharp elevation in H3K4me3 levels and permanently higher levels of H3K4me1, with variable fluctuations in H3K27ac and H3K27me3 levels.

We then measured mean CpG methylation levels of each region in mouse oocytes and sperm using data from Wang et al.(123) analyzed by the Smith Lab(106) to explore germline inheritance of allele-specific CpG methylation in imprinted-like regions (Table S3). Known imprinted regions exhibited anticipated oocyte- or sperm-specific CpG methylation, with the *Meg3* imprinted region having similar levels in oocyte and sperm as it acquires its imprinted status during development (Figure 4D). Differential CpG methylation between gametes was also observed in imprinted-like regions (Figure 4E); for instance, *Repin1* and *Errfi1 downstream* showed oocyte specificity. Displaying gamete and mESC epigenetic profiles of *Repin1* and *Errfi1 downstream* imprinted-like regions alongside those of oocyte-specific known imprinted regions *Mest* and *Plagl1* clearly showcased CpG hypermethylation in oocytes versus sperm and similar epigenetic alteration dynamics during *Dnmt1* inactivation and rescue in mESCs, especially for CpG methylation, H3K4me3, H3K4me1, and H3K9me3 (Figure S3F). While the distinction of gamete-specific CpG methylation in imprinted-like regions was generally less pronounced compared to known imprinted regions, they all exhibited substantial levels (> 35%) in at least one gamete (Figure 4D-E; Table S3). This suggests germline inheritance of CpG methylation profiles, albeit not in an allele-specific manner for all imprinted-like regions. Allele specificity may be acquired post-fertilization for some imprinted-like regions, as is the case for the *Meg3* imprinted region.

To further investigate regulation of imprinted-like regions by genomic imprinting mechanisms, we conducted enrichment analysis of ZFP57 DNA-binding motif (Figure 4F), revealing positive matches with 80% of imprinted-like regions and, predictably, with 100% of known imprinted regions (Figure 4G). Moreover, the permanent decrease of broad H3K9me3 detected in our model was strikingly mirrored in *Zfp57* KO mESCs(139) and *Setdb1* KO mESCs(140) for known imprinted regions (e.g. *H19*, *Peg13–Trappc9*, *Zrsr1–Commd1*) and imprinted-like regions (e.g. *Zfp13*, *Rnf216 upstream*, *Zf668*), with binding of ZFP57 and SETDB1 coinciding within these regions (Figure 4H). This was observed regardless of gamete-specific CpG methylation, as evidenced by the *Zfp668* imprinted-like region displaying nearly identical profiles in both gametes.

Finally, we examined significant gene expression dysregulation (adjusted p-value ≤ 0.05) in *Dnmt1**^INV^*** and *Dnmt1**^RES^***versus *Dnmt1**^CTL^*** for imprinted genes controlled by the selected 15 known imprinted regions (Figures S4A-B, Table S4) and for genes associated with imprinted-like regions (Figures S4C-D, Table S4). As expected, numerous imprinted genes showed significant permanent downregulation (e.g. *Igf2*, *Trappc9*, *Cdkn1c*, *Phlda2*) or upregulation (e.g. *Mest*, *Plagl1*, *H19*, *Zrsr1*), mostly exhibiting strong dysregulation (log_2_ fold change ≤ −0.6 or ≥ 0.6). Some genes associated with imprinted-like regions were also significantly permanently downregulated (*Zfp668*) or upregulated (*Repin1*, *Rnf216*, *Hcn2*, *Vsig10*), albeit with milder dysregulation. Nevertheless, 17 out of the 22 genes associated with imprinted-like regions were dysregulated in *Dnmt1**^INV^***, possibly suggesting a transcriptional impact linked to de-repression of their imprinted-like region. However, given that imprinted-like regions are located outside gene promoters (Table S3) and exhibit fluctuations in enhancer-associated marks H3K4me1 and H3K27ac, their regulatory target genes may be positioned at more distal locations. To directly assess their influence on gene expression regulation, a targeted de-repression approach, such as CRISPR-mediated demethylation, would be required.

To summarize, our results suggest that, beyond H3K9me3, other histone modifications, notably enhancer-associated H3K4me1, play a role in the heritability of DNA methylation at imprinted loci. Our analyses also led to the identification of 20 imprinted-like regions exhibiting epigenetic and regulatory features closely resembling those of known imprinted regions. Interestingly, three imprinted-like regions (i.e. *Spred2*, *Lipt1-Mitd1* and *Zfp668*) were also identified as such in mouse preimplantation embryos(159). While all imprinted-like regions had substantial DNA methylation levels in gametes, only some showed gamete specificity, with many displaying similar levels in both oocytes and sperm. Genomic imprinting establishment could thus be occurring in gametes for some imprinted-like loci whereas others may become imprinted during development.

### Permanent de-repression of MERVL and MT2 LTRs may be linked to gene transcript chimerism

DNA methylation participates in the repression of transposable elements(160). Loss of DNA methylation can lead to transposable element de-repression, which in turn can influence gene expression regulation(161) and induce genomic instability(162). LTRs and LINEs being significantly susceptible to permanent CpG hypomethylation during *Dnmt1* inactivation and rescue (Figure 3B) prompted us to determine whether this was accompanied by increased transcriptional activation, indicative of de-repression. Additionally, we sought to explore how their de-repression might impact the regulation of neighboring genes through the generation of chimeric gene transcripts(163).

We initially investigated whether specific LTR and LINE families showed distinct propensities for permanent CpG hypomethylation. The frequency of permanently hypomethylated CpGs within the most abundant LTR families (Figure 5A)—ERV1 (0.28), ERVK (0.33), ERVL (0.29), and ERVL-MaLR (0.26)—did not substantially deviate from the overall frequency in LTRs (0.31) (Figure 3A). The frequency in L1 LINEs (0.52) (Figure S5A), the most abundant LINE family, was identical to all LINEs combined (Figure 3A). Though, when computing the frequencies of permanently hypomethylated CpGs specifically in high-occupancy elements (i.e. genomic occurrences ≥ 500), the vast majority had strikingly higher frequencies (Figures 5A and S5A). We thus focused on high-occupancy elements to explore if their permanent CpG hypomethylation affected their transcription. Mean CpG methylation levels and total normalized RNA-seq counts of all loci combined were measured for each high-occupancy element across conditions. All high-occupancy L1 elements exhibited permanent mean CpG hypomethylation ≥ 10% (Figure S5B), but none showed major alterations in transcription (total normalized count log_2_ fold change > −1 and < 1 in *Dnmt1**^INV^***and *Dnmt1**^RES^*** versus *Dnmt1**^CTL^***) (Figure S5C). Turning toward high-occupancy LTR elements, MERVL-int and MT2_Mm emerged as the sole elements demonstrating two key characteristics: permanent mean CpG hypomethylation ≥ 10% (Figure 5B) and notable gradual increase in transcription throughout *Dnmt1* inactivation and rescue, culminating with a log_2_ fold change ≥ 1 in *Dnmt1**^RES^***versus *Dnmt1**^CTL^*** (Figure 5C). Given the evidence supporting the pivotal roles of MERVL-int and MT2_Mm in zygotic genome activation(164–166) and potency state transitions(167,168), we undertook a deeper analysis of locus-specific CpG methylation and transcription dynamics. Overall, MERVL-int and MT2_Mm showed highly significant decrease in locus-specific mean CpG methylation levels and increase in locus-specific transcript levels in *Dnmt1**^INV^***and *Dnmt1**^RES^*** compared to *Dnmt1**^CTL^***(Figures 5D-G; Table S5). When examining each locus, permanent but gradual increase in transcription occurred at loci exhibiting either temporary or permanent CpG hypomethylation (Figures 5H-I). Moreover, MERVL-int and MT2_Mm upregulation proximal to gene transcription start sites (TSS) (e.g. *Arap2*, *Cwc22*, *Zfp677*) suggests the generation of associated chimeric gene transcripts (Figure 5J).

**Figure 5:**
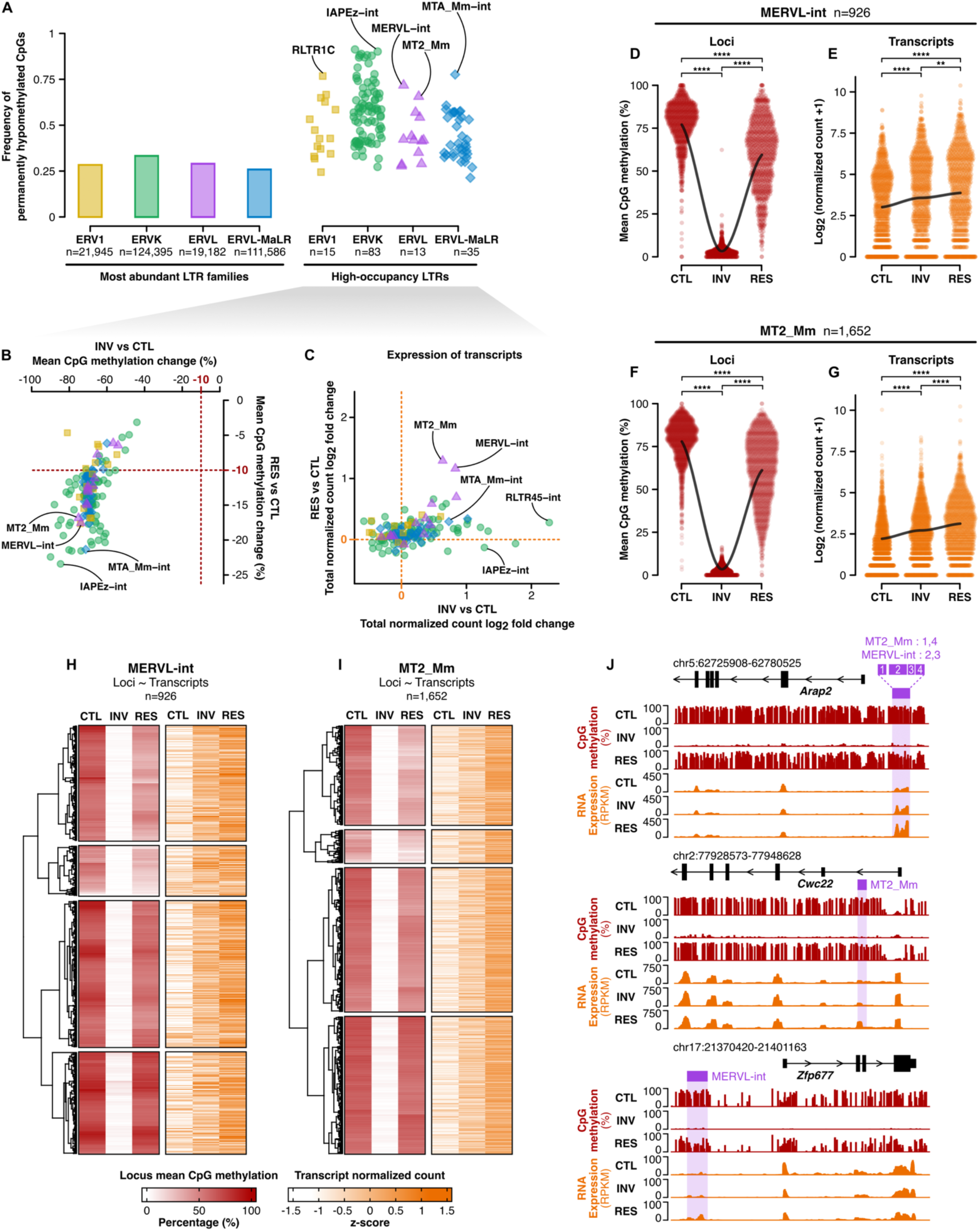
Permanent de-repression of MERVL and MT2 LTRs may be linked to gene transcript chimerism. **(A)** Frequency of permanently hypomethylated CpGs in the most abundant LTR families and in high-occupancy LTRs (genomic occurrences ≥ 500). **(B-C)** Change in mean CpG methylation levels (EM-seq) and log_2_ fold change of total RNA-seq normalized counts, respectively, of high-occupancy LTRs in *Dnmt1**^INV^*** and *Dnmt1**^RES^*** versus *Dnmt1**^CTL^*** (EM-seq: n=3, RNA-seq: n=2 per condition). **(D-E)** Pairwise comparisons of MERVL-int locus-specific mean CpG methylation levels and RNA-seq normalized counts in *Dnmt1**^CTL^***, *Dnmt1**^INV^*** and *Dnmt1**^RES^*** using the Wilcoxon rank sum test with Benjamini-Hochberg p-value adjustment. **(F-G)** Pairwise comparisons of MT2-Mm locus-specific mean CpG methylation levels and RNA-seq normalized counts in *Dnmt1**^CTL^***, *Dnmt1**^INV^*** and *Dnmt1**^RES^*** using the Wilcoxon rank sum test with Benjamini-Hochberg p-value adjustment. **(H-I)** Z-score normalization of locus-specific CpG methylation level and RNA-seq normalized count for MERVL-int and MT2_Mm. **(J)** Genomic signal tracks showing CpG methylation and RNA expression for MERVL-int and MT2-Mm loci proximal to transcription start sites. *See also Table S5*.

To sum up, high-occupancy LTRs and LINEs exhibited heightened susceptibility to permanent CpG hypomethylation during *Dnmt1* inactivation and rescue. MERVL-int and MT2_Mm LTRs stood out by displaying a progressive increase in transcriptional activation, even at loci where loss of DNA methylation was only temporary. This could reflect a form of epigenetic memory, resulting from changes in chromatin architecture or in the binding of regulatory proteins that persist even after DNA methylation is restored. Additionally, loss of DNA methylation could trigger feedback loops with distal regulatory elements that sustain transcriptional activation, even after the initial trigger is removed. Moreover, in some cases, the resilience of transposable element activation might arise from selective cellular advantages, such as through the generation of chimeric gene transcripts that can forge novel functional outcomes.

### Gene expression dysregulation and genomic instability

We next assessed the broader molecular impact of transient DNMT1 loss in mESCs, extending beyond altered epigenomic landscapes. Specifically, we measured changes in gene expression, which can directly result from epigenomic alterations, and examined genomic stability, including the presence of DNA damage and the length of telomeres. DNMT1 contributes to preventing the accumulation of DNA damage by restoring methylation during DNA repair(34), and telomere length control has been linked to DNA methylation regulation(169–171).

We first examined significantly differentially expressed genes (adjusted p-value ≤ 0.05) in *Dnmt1**^INV^*** and *Dnmt1**^RES^*** conditions versus *Dnmt1**^CTL^***, focusing on genes exhibiting dysregulation of considerable magnitude (log_2_ fold change ≤ −0.6 or ≥ 0.6). Thousands of genes were dysregulated in *Dnmt1**^INV^*** (downregulated: n=2,714; upregulated; n=3,204) (Figure 6A; Table S6), whereas in *Dnmt1**^RES^***, the number was notably reduced but still in the hundreds (downregulated: n=460; upregulated; n=278) (Figure 6B; Table S6). To reveal patterns of gene expression dysregulation, we categorized genes based on their downregulation, upregulation, or stable expression in *Dnmt1**^INV^***and *Dnmt1**^RES^*** (Figures 6C and S6A). Interestingly, *Dnmt3a* and *Dnmt3b* exhibited temporary downregulation, whereas *Dnmt3l* demonstrated dynamic up- and downregulation. Conversely, other key players in DNA methylation such as *Uhrf1*, *Tet1-3*, *Zfp57*, and *Trim28*, maintained stable expression. *Kdm1a*, known for gene repression via H3K4 demethylation, was notably temporarily downregulated, possibly contributing to the marked H3K4me3 increase in *Dnmt1**^INV^***(Figure 2A). Dysregulation extended to cell cycle regulators, with *Myc*, *Ccnd1-2*, and *Chek2* exhibiting temporary downregulation, and *Ccng2* showing latent upregulation. Additionally, cellular stress and DNA damage response genes were affected, with *Ndrg1* and *Ddit4* showing latent upregulation, and *Ndrg1* showing dynamic down- and upregulation. Potency state regulators were also dysregulated, including temporary downregulation of *Dppa3* and permanent downregulation of *Zscan4a-f* and *Tcstv3*, with *Zscan4a-f* and *Tcstv3* being specifically recognized for their role in telomere elongation(172,173). Notably, an increase and spreading of repressive H3K9me3 across the promoters and gene bodies of *Zscan4a* and *Zscan4d* was evident in *Dnmt1**^INV^*** and *Dnmt1**^RES^*** (Figure S6B). Moreover, permanent upregulation of *Xlr3a-b* and *Xlr4a-b*, part of an X-linked cluster known for tissue-specific imprinted gene expression in mice(174), and of *Uba1y*, *Eif2s3y*, *Uty*, and *Ddx3y* within a Y-linked cluster were of particular interest given the significant susceptibility of sex chromosomes to permanent CpG hypomethylation (Figure 3B). In fact, promoters of *Xlr3a*, *Xlr4a*, *Eif2s3y* and *Uty* showed permanent CpG hypomethylation and increase of permissive H3K4me3 and H3K27ac in *Dnmt1**^INV^*** and *Dnmt1**^RES^*** (Figure S6C). Despite extensive and dynamic changes in gene expression, key pluripotency markers (*Pou5f1*, *Sox2*, *Nanog*) remained stably expressed during *Dnmt1* inactivation and rescue (Figure 6C), suggesting that mESC pluripotency and differentiation potential were generally preserved.

**Figure 6:**
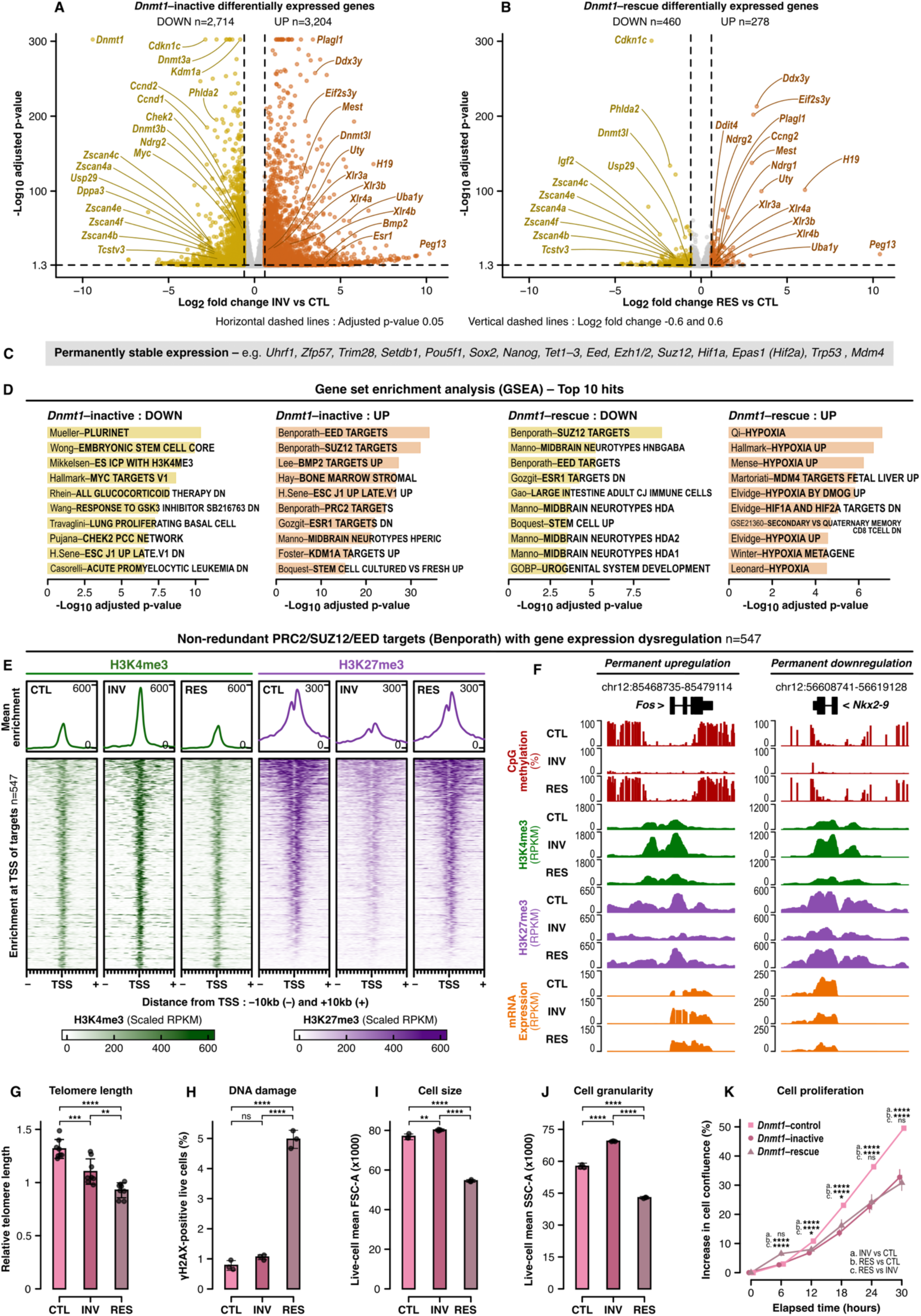
Gene expression dysregulation, genomic instability, and altered cellular processes. **(A-B)** Differentially expressed genes in *Dnmt1**^INV^*** and *Dnmt1**^RES^*** versus *Dnmt1**^CTL^*** determined by an adjusted p-value ≤ 0.05 and log_2_ fold change ≤ −0.6 or ≥ 0.6 using mRNA sequencing (n=3 per condition) analyzed with the DESeq2 software. **(C)** Notable genes with stable expression in *Dnmt1**^INV^*** and *Dnmt1**^RES^*** versus *Dnmt1**^CTL^***. **(D)** Gene set enrichment analysis for down- and upregulated genes in *Dnmt1**^INV^*** and *Dnmt1**^RES^***. **(E)** Signal enrichment of H3K4me3 and H3K27me3 in *Dnmt1**^CTL^***, *Dnmt1**^INV^*** and *Dnmt1**^RES^*** at transcription start sites of PRC2/SUZ12/EED non-redundant targets exhibiting gene expression dysregulation in *Dnmt1**^INV^*** and/or *Dnmt1**^RES^*** identified in D. RPKM values were scaled between 0 and 0.99 quantiles to mitigate the presence of outliers. **(F)** Genomic signal tracks showing CpG methylation, H3K4me3, H3K27me3 and mRNA expression for PRC2/SUZ12/EED targets *Fos* and *Nkx2-9* in *Dnmt1**^CTL^***, *Dnmt1**^INV^*** and *Dnmt1**^RES^***. **(G)** Pairwise comparisons of relative telomere length in *Dnmt1**^CTL^***, *Dnmt1**^INV^*** and *Dnmt1**^RES^*** measured by qPCR and the delta-delta Ct method, with normalization to reference single-copy gene *Rplp0* and relativization to reference DNA from R1 mESCs (n=8 per condition). **(H-J)** Pairwise comparisons of the percentage of γH2AX-positive live cells (DNA damage), live-cell mean forward scatter area (FSC-A; cell size), live-cell mean side scatter area (SSC-A; cell granularity i.e. internal complexity), respectively, in *Dnmt1**^CTL^***, *Dnmt1**^INV^*** and *Dnmt1**^RES^*** measured by FACS (n=3 per condition). **(K)** Pairwise comparisons of cell proliferation rate for *Dnmt1**^CTL^***, *Dnmt1**^INV^*** and *Dnmt1**^RES^*** measured by the increase in confluency over 30 hours (n=3 per condition). All pairwise comparisons were conducted using one-way ANOVA with Tukey’s HSD test with p-value adjustment. *See also Tables S6 and S7 and Figures S6 and S7*.

We then proceeded with gene set enrichment analysis (GSEA) of down- and upregulated genes in *Dnmt1**^INV^*** and *Dnmt1**^RES^***, selecting their top 10 significantly enriched gene sets (Figure 6D). EED(175), SUZ12(175), PRC2(175), and KDM1A(176) targets stood out among upregulated gene sets in *Dnmt1**^INV^***, and, interestingly, EED and SUZ12 targets were conversely enriched in *Dnmt1**^RES^*** downregulated genes. PRC2, comprising SUZ12 and EED among its components, mediates the deposition of H3K27me3 in bivalent developmental gene promoters(177). Nearly all promoters of dysregulated PRC2/SUZ12/EED target genes(175) (non-redundant: n=547) exhibited a simultaneous sharp surge in H3K4me3 and decline in H3K27me3 centered at their TSS in *Dnmt1**^INV^***, with levels returning to baseline in *Dnmt1**^RES^*** (Figure 6E). This temporary transition to an active chromatin state occurred independently of changes in CpG methylation levels within TSS regions, which remained consistently very low across all conditions (median levels < 5.5%), and independently of gene expression dysregulation patterns (Figure S6D; Table S7), as exemplified by epigenetic profiles of *Fos* and *Nkx2-9* (Figure 6F). Additional compelling insights emerged from our GSEA analysis. Pluripotency network(178), embryonic stem cell core(179), MYC targets(136), and CHEK2 network(180) gene sets were enriched among *Dnmt1**^INV^***downregulated genes, while *Dnmt1**^RES^*** upregulated genes showed predominant enrichment in hypoxia gene sets(136,181–185), suggesting disturbances cell cycle regulation and an increase in cellular stress. In fact, throughout *Dnmt1* inactivation and rescue, we detected signs of genomic instability, indicative of cellular stress. We observed a progressive shortening of telomere length (Figure 6G) and a latent increase in yH2AX (Figure 6H), a marker for DNA double-strand breaks.

In summary, while *Dnmt1* rescue restored the expression of most genes that had been dysregulated following its inactivation, not all expression levels returned to baseline, with the rescue also resulting in additional gene expression dysregulation. Dysregulation of cell cycle regulators, downregulation of telomere elongation factors and upregulation of cellular stress and DNA damage response genes were notably detected, corresponding with *Dnmt1* inactivation and rescue leading to shortened telomeres and increased DNA damage. These results highlight the far-reaching molecular consequences of lacking continuous DNA methylation maintenance, extending beyond altered epigenomic landscapes by disrupting gene expression levels and inducing genomic instability.

### Morphological and functional cellular responses

ESCs are characterized by their high cellular plasticity, which grants them resilience to significant epigenetic perturbations(36,76–78). Still, we questioned whether the molecular alterations caused by transient *Dnmt1* inactivation would manifest in cellular responses. To address this, we assessed mESC morphology (size, internal complexity) and functional properties, including proliferation rate and differentiation potential toward germ layer cell types (endoderm, mesoderm, ectoderm). The impact on gene expression patterns in germ layer cells was also examined.

Dynamic changes in cell size and internal complexity (i.e. granularity) were observed throughout *Dnmt1* inactivation and rescue in mESCs, increasing in *Dnmt1**^INV^*** and decreasing in *Dnmt1**^RES^*** (Figures 6I-J and S7A), and cell proliferation rate (Figure 6K) was reduced in both *Dnmt1**^INV^*** and *Dnmt1**^RES^***. These results reflect altered cell cycle dynamics, consistent with DNMT1’s critical role during cell division(29–32). The molecular and cellular alterations in mESCs prompted us to investigate if temporary inactivation of *Dnmt1* would have a lasting impact on their differentiation. To verify this, *Dnmt1**^CTL^*** and *Dnmt1**^RES^*** mESCs were differentiated into endoderm, mesoderm, and ectoderm monolayers (Figure 7A) which were sorted based on lineage-specific cell surface markers. Similar percentages of marker-positive cells were obtained in *Dnmt1**^CTL^*** and *Dnmt1**^RES^***derived germ layer lineages (Figure S7B), and principal component analysis of gene expression showed comparable separation of mESCs and differentiated cells for both *Dnmt1**^RES^**Dnmt1**^CTL^*** conditions(Figure S7C). *Dnmt1**^CTL^*** and *Dnmt1**^RES^*** ectoderm inductions were similarly successful; compared to mESCs, pluripotency markers were all markedly lower, and differentiation markers were all markedly higher, whereas some, mostly minor, differences in marker levels between *Dnmt1**^CTL^*** and *Dnmt1**^RES^*** inductions were noticeable for endoderm and mesoderm (Figure S7D-F). Overall, transient inactivation of *Dnmt1* therefore did not appear to have a major impact on the differentiation potential of mESCs. However, differential gene expression analysis in *Dnmt1**^RES^***versus *Dnmt1**^CTL^*** derived germ layers revealed hundreds of dysregulated genes (Figure 7B; Table S6), compellingly showing enrichment for PRC2(175), SUZ12(175) and EED(175) targets (Figure 7C) as was observed in mESCs (Figure 6D). Among *Dnmt1**^RES^***cell types, endoderm had the highest number of dysregulated genes, followed by mESCs, mesoderm, and ectoderm (Figure 7D). Down- and upregulated genes in *Dnmt1**^RES^*** cell types were primarily cell type-specific, though some were shared among multiple *Dnmt1**^RES^*** cell types such as *Dnmt3l* downregulated in mESCs, endoderm and mesoderm, and *Sall3* upregulated in all germ layers (Figures 7E-F).

**Figure 7:**
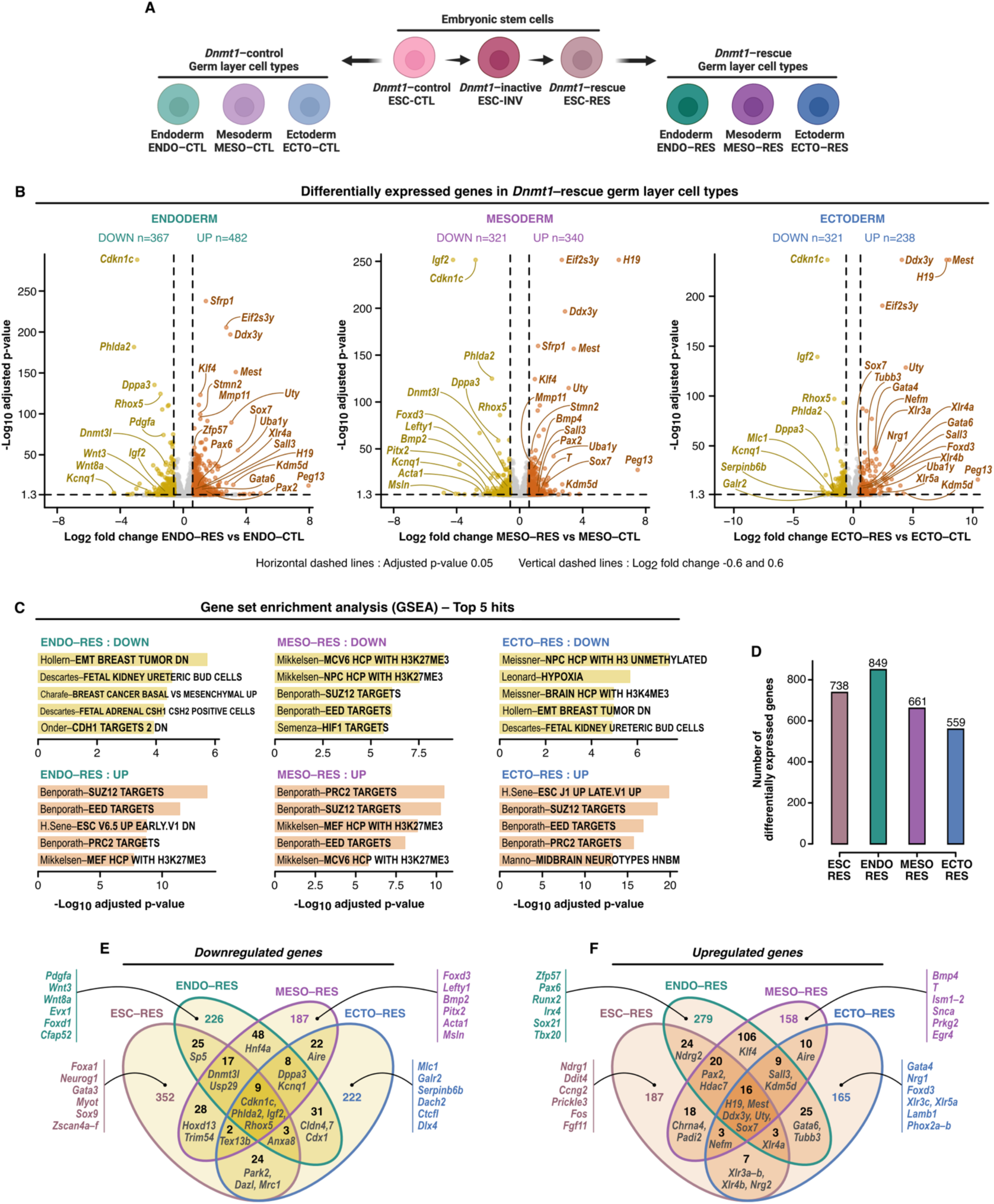
Lasting effects on gene expression patterns upon differentiation to embryonic germ layers. **(A)** Schematic diagram depicting *Dnmt1**^CTL^*** and *Dnmt1**^RES^***differentiation into germ layer monolayers (n=3 per differentiation). Made with Biorender.com. **(B)** Differential gene expression in *Dnmt1**^RES^***versus *Dnmt1**^CTL^*** endoderm, mesoderm and ectoderm determined by an adjusted p-value ≤ 0.05 and log_2_ fold change ≤ −0.6 or ≥ 0.6 using mRNA sequencing (n=3 per differentiation) analyzed with the DESeq2 software. **(C)** Gene set enrichment analysis for down- and upregulated genes in *Dnmt1**^RES^*** germ layers. **(D)** Total number of differentially expressed genes in *Dnmt1**^RES^*** cell types. **(E-F)** Overlap of downregulated and upregulated genes among *Dnmt1**^RES^***cell types. *See also Table S6 and Figure S7*.

Taken together, these findings demonstrate that despite their high cellular plasticity, mESCs exhibited morphological and functional responses to transient loss of DNMT1 activity, including reduced cell size, internal complexity and proliferation rate as well as altered gene expression profiles upon their differentiation toward germ layers. Dysregulation of gene expression in germ layers was mostly lineage-specific, illustrating how the impact of epigenetic perturbations can be cell type-specific, thus resulting in phenotypic variation depending on the affected cells(20–22). However, stemness properties remained well-preserved following *Dnmt1* inactivation and rescue in mESCs: pluripotency marker genes were stably expressed, mESCs retained the ability to proliferate indefinitely, albeit at a slower rate, and the potential of mESCs for differentiation toward germ layers was generally conserved.

### Delaying *Dnmt1* rescue until after differentiation initiation worsens molecular and cellular outcomes

We demonstrated that rescuing DNMT1 activity was insufficient to fully counteract the effects of its loss in mESCs, inducing permanent and post-rescue molecular and cellular responses as well as lasting effects upon differentiation toward ectoderm, mesoderm and endoderm germ layers. Next, we hypothesized that delaying *Dnmt1* rescue after differentiation initiation would lead to worse outcomes, as differentiated cells are typically less resilient to epigenetic disturbances due to cellular plasticity decreasing during differentiation(186,187). We tested this hypothesis using embryoid bodies which are three-dimensional (3D) spheroid aggregates of ESCs that spontaneously differentiate into early developmental cell types, enabling longitudinal studies of certain aspects of developmental processes(188,189).

Embryoid bodies were derived from *Dnmt1*–control mESCs, *Dnmt1*–rescue mESCs, as well as from *Dnmt1*–inactive mESCs for which *Dnmt1* inactivation was maintained throughout the 10 days of embryoid body formation. On day 5, *Dnmt1* was reactivated in a subset of *Dnmt1*–inactive embryoid bodies for post-differentiation rescue of *Dnmt1*. Four experimental conditions were thus included during embryoid body formation: *Dnmt1*–control, *Dnmt1*–inactive, *Dnmt1*–rescue and post-differentiation *Dnmt1*–rescue (Figure 8A). Successful inactivation and post-differentiation rescue of *Dnmt1* in embryoid bodies were confirmed by Western blot (Figure S8A). Throughout embryoid body formation, morphological and molecular analyses were conducted. Daily bright-field microscopy images were taken of embryoid bodies for measuring their size and circularity, and mRNA sequencing was performed on days 5 and 10.

**Figure 8:**
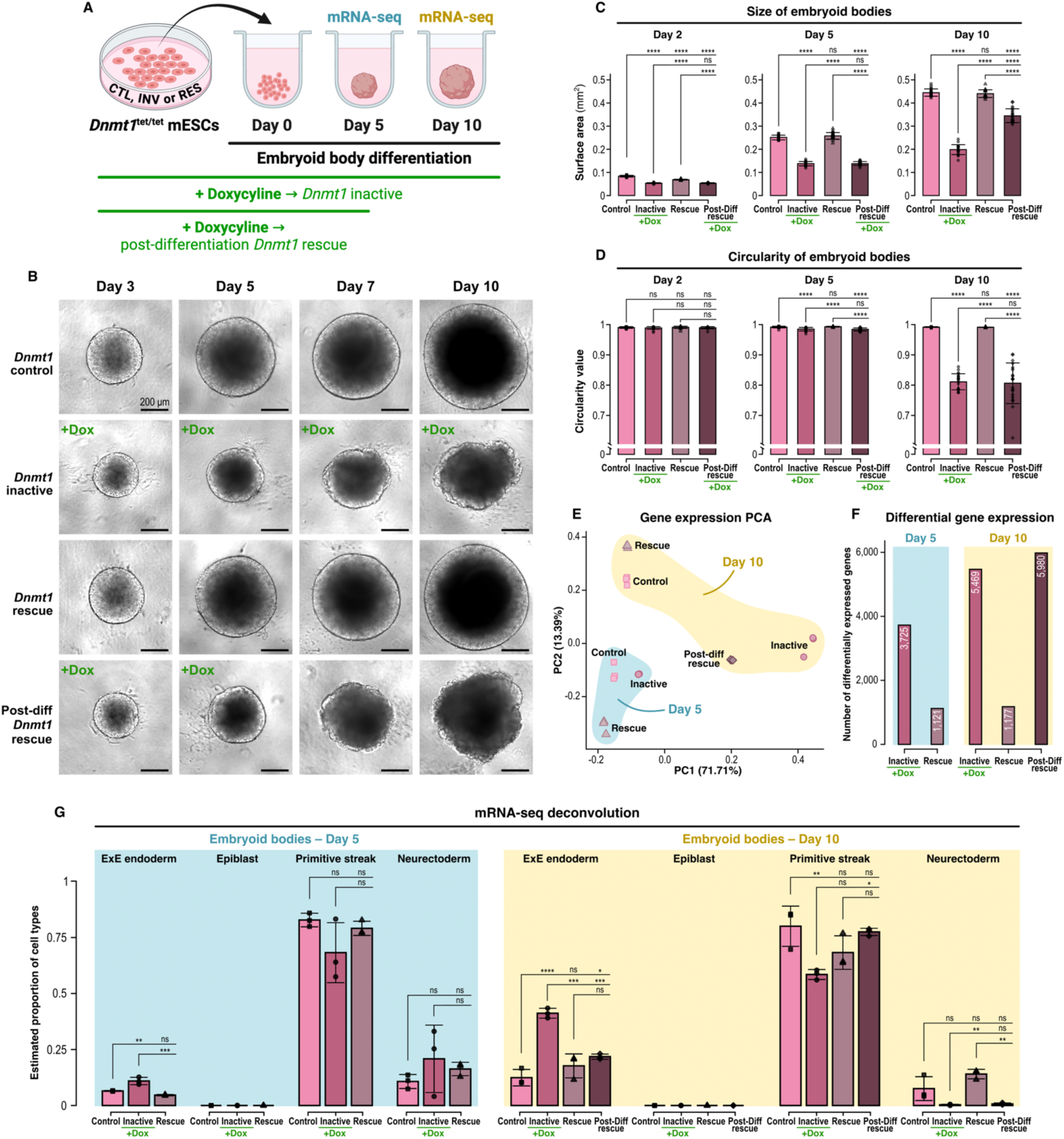
Delaying *Dnmt1* rescue until after differentiation initiation worsens molecular and cellular outcomes. **(A)** Schematic diagram depicting embryoid body formation including four experimental conditions: *Dnmt1*–control, *Dnmt1*– inactive, *Dnmt1*–rescue and post-differentiation *Dnmt1*–rescue. Embryoid bodies (n=192 per condition) were derived from the same *Dnmt1*^tet/tet^ mESC culture. Made with Biorender.com. **(B)** Size and **(C)** circularity of embryoid bodies on days 2, 5 and 10 determined by measuring their surface area and perimeter on bright-field microscopy images using the ImageJ software. Daily images were taken of the same 20 embryoid bodies per condition. Surface area was used as an indicator of size and circularity was calculated with the formula 4π × Area/Perimeter². **(D)** Bright-field microscopy images of one embryoid body per condition taken on day 3, 5, 7 and 10. **(E)** Principal component analysis of gene expression measured on day 5 and day 10 by mRNA sequencing (n=3 per condition with pooled embryoid bodies). A greater divergence in gene expression profiles is represented by the distance between samples along the x-axis (PC1), compared to the y-axis (PC2). **(F)** Number of differentially expressed genes in *Dnmt1*–inactive, *Dnmt1*–rescue and post–differentiation *Dnmt1*–rescue embryoid bodies versus *Dnmt1*–control embryoid bodies on day 5 and day 10 determined by an adjusted p-value ≤ 0.05 and log2 fold change ≤ −0.6 or ≥ 0.6 using mRNA sequencing analyzed with the DESeq2 software. **(G)** Estimated proportion of extraembryonic (ExE) endoderm, epiblast (pluripotent), primitive streak (mesendoderm) and neurectoderm cell types in embryoid bodies on days 5 and 10 determined by deconvoluting raw counts using single-cell data from mouse gastrulation embryos (embryonic day 7.75). All pairwise comparisons were conducted using one-way ANOVA with Tukey’s HSD test and p-value adjustment. *See also Figure S8*.

On day 2 of embryoid body formation, *Dnmt1*–rescue embryoid bodies were slightly smaller than *Dnmt1*–control embryoid bodies but rapidly caught up in size, whereas *Dnmt1*–inactive embryoid bodies remained substantially smaller throughout all 10 days of embryoid body formation (Figures 8B-C and S8B). Rescuing *Dnmt1* on day 5 in *Dnmt1*–inactive embryoid bodies triggered a marked, progressive increase in growth; by day 10, post-differentiation *Dnmt1*–rescue embryoid bodies were substantially larger than *Dnmt1*–inactive embryoid bodies, though still smaller than both *Dnmt1*–control and *Dnmt1*–rescue embryoid bodies (Figures 8B-C and S8B). However, post-differentiation rescue of *Dnmt1* could not prevent the eventual loss of embryoid body circularity resulting from *Dnmt1* inactivation during early embryoid body formation. Despite being reasonably circular up to day 5, *Dnmt1*–inactive and post-differentiation *Dnmt1*–rescue embryoid bodies progressively developed highly irregular shapes, while *Dnmt1*–control and *Dnmt1*–rescue embryoid bodies consistently maintained nearly perfect circularity (Figures 8B, 8D and S8C).

We then performed gene expression analysis on days 5 and 10 of embryoid body formation, revealing extensive alterations in *Dnmt1*-inactive embryoid bodies that worsened over time. Principal component analysis showed that their gene expression profiles progressively diverged from controls between days 5 and 10 (Figure 8E), with the number of differentially expressed genes increasing from 3,725 on day 5 to 5,469 on day 10 (Figure 8F). Post-differentiation *Dnmt1*-rescue embryoid bodies exhibited a higher number of significantly dysregulated genes than *Dnmt1*-inactive embryoid bodies on day 10 (Figure 8F), but their overall gene expression profiles were more aligned with controls (Figure 8E). Although *Dnmt1*–rescue embryoid bodies displayed similar growth and morphology compared to controls, gene expression was substantially affected, albeit to a lesser extent than in the *Dnmt1*-inactive and post-differentiation *Dnmt1*-rescue conditions (Figures 8E-F). Principal component analysis revealed distinct separation between the gene expression profiles of *Dnmt1*-rescue and *Dnmt1*-control embryoid bodies on days 5 and 10 of their formation (Figure 8E), with over 1,100 significantly differentially expressed genes identified at each time point (Figure 8F). Notably, the number of differentially expressed genes was considerably higher in embryoid bodies than in monolayer germ layers derived from *Dnmt1*–rescue mESCs (Figure 7D), likely due to reduced cellular plasticity in 3D versus monolayer cell cultures(190). Monolayer cell cultures often exhibit greater cellular plasticity than 3D cultures because they lack the spatial and mechanical constraints present in a 3D environment, allowing cells to more freely adapt their shape, interactions, and gene expression. Conversely, 3D cultures mimic in vivo conditions more closely, promoting cell polarity, differentiation, and tissue-specific behaviors that restrict plasticity.

Next, we deconvoluted gene expression profiles(147,148) using single-cell RNA-seq data from mouse gastrulation embryos(146) to estimate the proportions of inner cell mass-derived early developmental cell types within embryoid bodies across experimental conditions and time points (Figure 8G). Specifically, we assessed the proportion of extraembryonic (ExE) endoderm cells, epiblast cells (pluripotent precursors to embryonic germ layers), primitive streak cells (precursors to mesoderm and endoderm lineages), and neurectoderm cells (precursors to neural tissues). As expected, embryoid bodies across all experimental conditions showed no proportion of epiblast cells on days 5 and 10. Cell type proportions in *Dnmt1*–rescue embryoid bodies were similar to those in controls at both time points, while *Dnmt1*–inactive and post-differentiation *Dnmt1*–rescue embryoid bodies exhibited significant differences. Most notable was the higher proportion of ExE endoderm cells in *Dnmt1*–inactive embryoid bodies that continued to increase over time, with post-differentiation rescue of *Dnmt1* partially preventing this cell type proportion shift toward ExE endoderm cells. This observation aligns with findings from a previous study showing that DNA methylation is essential for the differentiation of embryonic lineages but dispensable for extraembryonic lineages; triple DNMT knockout mESCs exhibited progressive cell death during their differentiation into epiblast-derived cells but maintained viability when differentiating into extraembryonic cells, both in vitro and in vivo(191).

Overall, this experiment demonstrated that *Dnmt1* inactivation in mESCs and throughout embryoid body differentiation caused severely stunted growth, abnormal morphology and gene expression alterations, accompanied by a striking shift in cell type proportions, with a more prominent representation of extraembryonic compared to embryonic cell types. Rescuing *Dnmt1* midway through embryoid body formation partially restored growth and cell type proportions and mitigated the severity of gene expression alterations but failed to prevent severe morphological abnormalities. In contrast, rescuing *Dnmt1* in mESCs prior to embryoid body formation supported normal growth, morphology and cell type proportions and led to only moderate differences in gene expression, but the impact on gene expression was more pronounced than in monolayer differentiated cells. These findings illustrate the escalating impact of an epigenetic perturbation as cellular plasticity decreases and demonstrate the disadvantages of a delayed epigenetic rescue intervention, underscoring the importance of early intervention for better developmental outcomes.

## DISCUSSION

In this study, *Dnmt1* inactivation and rescue in mESCs had a wide-ranging molecular impact, leading to DNA methylation and histone modification alterations, gene expression dysregulation, genomic instability and transposable element de-repression. Transient *Dnmt1* inactivation also induced morphological and functional responses in mESCs and had lasting effects upon their differentiation, with post-differentiation rescue of *Dnmt1* worsening outcomes. Moreover, transient *Dnmt1* inactivation in mESCs enabled the discovery of 20 regions with imprinted-like signatures.

### Genomic imprinting mechanisms and discovery of novel imprinted regions

Mouse ESC and embryo models remain invaluable tools for deciphering genomic imprinting mechanisms and uncovering novel imprinted regions through disruption of genomic imprinting maintenance machinery(36,37,139,159). However, characterization of imprinted regions is highly complex due to diverse regulatory factors and epigenetic modifications, often defying a robust universal mechanistic paradigm.

While DNA methylation and H3K9me3 are extensively investigated in genomic imprinting, cellular and animal mouse studies have implicated additional histone modifications. Imprinted (i.e. repressed) alleles, co-marked by DNA methylation and H3K9me3(84,107,159,192), may occasionally also display H3K27me3(84,192), whereas H3K4me2/3(84,192,193) and H3K27ac(194) characterize permissive alleles, which can exhibit bivalency with concurrent H3K27me3(84,192,193). This study was distinctive for simultaneously profiling H3K4me3, H3K4me1, H3K27ac and H3K27me3, alongside customary DNA methylation and H3K9me3, offering an unprecedented perspective on the complex and divergent nature of imprinting mechanisms. While de-repression of imprinted regions following *Dnmt1* inactivation, marked by DNA hypomethylation and decreased H3K9me3, was consistently accompanied by increases in H3K4me3 and H3K4me1, fluctuations in H3K27ac and H3K27me3 varied across different imprinted regions, illustrating the divergent nature of imprinting mechanisms. In certain instances, increased H3K4me3 overlapped with increased H3K4me1, and increased H3K27ac overlapped with H3K27me3—an intriguing observation given that these modification pairs are mutually exclusive as they occur on the same amino acid residue. This could suggest that imprinted regions harbor asymmetrical nucleosome modifications(195) or exhibit cellular heterogeneity in their epigenetic profiles. Additionally, these pairs of mutually exclusive histone modifications might reside on different alleles. De-repression of the imprinted allele could induce epigenetic alterations on the opposing allele through transvection effects, a form of epigenetic crosstalk between alleles(196–199). Further investigation of these potential mechanisms would benefit from the use of mESCs with parental single-nucleotide polymorphisms, enabling allele-specific profiling of epigenetic marks and chromatin interactions. Moreover, the role of H3K4me1 in genomic imprinting has remained unexplored until now, but its correlation with enhancer activity(200–203) imputes relevance for permissiveness of imprinted loci, a notion supported by the increased H3K4me1 at de-repressed imprinted regions detected this study. H3K4me1 can help distinguish promoter versus enhancer activity, and active versus poised states, valuable nuances for elucidating how imprinted regions regulate gene expression.

Epigenetic analysis of imprinted regions during *Dnmt1* inactivation and rescue in mESCs uncovered potential novel imprinted regions. These imprinted-like regions exhibited remarkably similar epigenetic responses to transient *Dnmt1* inactivation as those observed in known imprinted regions. Notably, three of the imprinted-like regions identified in this study—*Spred2*, *Lipt1-Mitd1* and *Zfp668*—have also been characterized as imprinted-like in mouse preimplantation embryos(159). Imprinted regions play a crucial role in development by regulating the expression of genes that are predominantly essential for developmental processes(154–156). In humans, *LIPT1* is the causative gene for lipoyltransferase 1 deficiency, a rare inborn metabolism disorder with which *MITD1* is also associated(204–207), and *SPRED2* is a causative gene for Noonan syndrome 14(208), a severe developmental disorder. Additional imprinted-like regions discovered in this study may also be linked to genes involved in human diseases, such as *REPIN1* linked to non-alcoholic fatty liver disease(209) and polycystic ovary syndrome(210), *HCN2* to epilepsy(211–214), *RNF216* to cerebellar ataxia-hypogonadism syndrome(215,216) and *VSIG10* to COVID-19(217). In this study, these genes became dysregulated upon *Dnmt1* inactivation, when imprinted-like regions were de-repressed, suggesting that their regulation may possibly be influenced by their associated imprinted-like region. Therefore, future work exploring the developmental importance of these imprinted-like regions could help unveil novel associations with human diseases.

### Wide-ranging molecular and cellular influence of DNMT1

Beyond genomic imprinting maintenance, DNMT1 maintains global DNA methylation profiles(29–32) and contributes to preserving genomic stability(34) during cell division. Through non-catalytic mechanisms(55–58), it can also influence broader epigenomic networks(59,60), including chromatin conformation, histone modifications and gene expression.

In the S phase of the cell cycle, DNMT1 methylates newly synthesized DNA(29–32) through its interaction with proliferating cell nuclear antigen (PCNA)(218,219) and restores methylation following DNA repair(34). Stressful conditions such as viral infection(220), inflammation(221) or DNA hypomethylation due to DNMT1 depletion(222) can induce cell cycle arrest in G1, preventing DNA damage accumulation in S phase. Progression to S phase despite stress can increase DNA damage, triggering the G2/M checkpoint for G2 arrest(223,224). In this study, *Dnmt1* inactivation in mESCs led to downregulation of *Ccnd1/2*, essential for G1/S transition(225–228), whereas *Ccng2*, implicated in G1 and G2 arrest(229–232), was upregulated after *Dnmt1* rescue alongside increased DNA damage. Permanent reduction of mESC proliferation rate throughout inactivation and rescue of *Dnmt1* could therefore be linked to increased cell cycle arrest, in G1 for *Dnmt1**^INV^*** due to drastic DNA methylation and histone modification alterations and in either G1 or G2 for *Dnmt1**^RES^***. Less severe epigenomic changes in *Dnmt1**^RES^*** could have allowed some cells to escape G1 arrest, culminating in DNA damage accumulation and G2/M checkpoint activation. Another potential trigger of cell cycle arrest in *Dnmt1**^INV^***and *Dnmt1**^RES^*** was telomere shortening(233–235), which may have occurred due to permanent downregulation of *Zscan4a-f(172)* and *Tcstv3(173)*, required for telomere elongation in mESCs.

Moreover, *Zscan4a-f* and *Tcstv3* are part of a mouse 2-cell embryo (2C) gene network activated in rare mESC subpopulations as they cycle in and out of a 2C-like totipotent state(167,168). This 2C-like totipotent state is further characterized by decreased global DNA methylation levels, absence of pluripotency marker OCT4/POU5F1, de-repression of MERVL and MT2 LTRs and generation of MERVL- and MT2-derived chimeric gene transcripts(167,168). This study showed that *Dnmt1* inactivation and rescue in mESCs led to permanent reduction of global DNA methylation and permanently de-repressed MERVL and MT2 LTRs with evidence of associated gene transcript chimerism but did not affect *Oct4*/*Pou5f1* expression levels. This could indicate atypical potency state transitions in *Dnmt1**^RES^*** mESCs, potentially compromising their proliferation rate and other cellular properties, such as their morphological features. In fact, *Dnmt1**^RES^*** mESCs exhibited reduced cell size and internal complexity. Conversely, Eckersley-Maslin et al. did not observe MERVL upregulation upon *Dnmt1* repression in mESCs(168). However, they repressed *Dnmt1* for only three days, after which mean whole-genome and MERVL-specific DNA methylation were both reduced to ∼30%, in contrast to six days of *Dnmt1* repression in this study, where mean levels were reduced to ∼3.5% and ∼3.2%, respectively. Absence of DNMT1 for just three days was likely insufficient to trigger the same magnitude of effects observed in this study, where DNMT1 was absent for twice as long, highlighting how the duration of an epigenetic perturbation significantly influences the severity of the resulting outcomes.

The impact of transient *Dnmt1* inactivation on cellular functions was further evidenced by altered gene expression profiles in germ layers derived from *Dnmt1**^RES^*** mESCs. Lineage-specific and pan-lineage gene expression alterations were observed, with endoderm and mesoderm being more severely affected than ectoderm. Downregulation of *Dnmt3l* in *Dnmt1**^RES^*** mESCs persisting only in endoderm and mesoderm could have instigated deviations in their differentiation trajectories, as DNMT3L is a crucial co-factor for DNMT3A-mediated *de novo* DNA methylation(236,237), essential for orchestrating mammalian embryonic development(238). Upregulation of *Sall3* in all *Dnmt1**^RES^*** germ layers may have also contributed to lineage-specific effects. In human induced pluripotent stem cells, SALL3 serves as a predictive marker for differentiation propensity, positively correlating with ectoderm and negatively with endoderm/mesoderm, mechanistically via modulation of DNMT3B function(239), another vital *de novo* DNA methyltransferase(238). Gene expression regulation may have been supported by *Sall3* upregulation in *Dnmt1**^RES^***ectoderm, while being hindered in *Dnmt1**^RES^*** endoderm and mesoderm. Additionally, sex chromosome genes were notably upregulated among *Dnmt1**^RES^*** cell types: X-linked *Xlr3a-b/4b* in mESCs and ectoderm, *Xlr4a* in mESCs, endoderm, and ectoderm, and *Xlr3c/5a* in ectoderm, as well as Y-linked *Uty*, *Ddx3y*, and *Eif2s3y* in all cell types, and *Kdm5d* in all three germ layers. The upregulation of X- and Y-linked genes aligns with sex chromosomes being significantly susceptible to permanent DNA hypomethylation in mESCs following transient *Dnmt1* inactivation. Collectively, these findings relate to how epigenetic perturbations can often have differential consequences based on cell type(20–22), with further distinctions due to sex-specific effects(11,23).

### Repercussions of epigenetic perturbations during embryonic development

In this study, the severe growth delay and morphological abnormalities of mouse embryoid bodies resulting from DNMT1 deficiency bared strong resemblance to the phenotypes of DNMT1-deficient mouse embryos, which die by mid-gestation, stunted and deformed(61). Despite partial growth recovery, the persistent severe malformation of mouse embryoid bodies after rescuing DNMT1 post-differentiation suggests that such delayed intervention in vivo, at gastrulation or later developmental stages, would likely fail to avert the lethal embryonic outcomes of DNMT1 deficiency. An earlier rescue intervention during stages of greater cellular plasticity, such as the blastocyst stage where pluripotent stem cells are prominent, may better support embryo viability, as evidenced by the normal growth and morphology of embryoid bodies in which DNMT1 was rescued in mESCs before their differentiation. However, epigenetic perturbations can result in drastically worse effects in vivo than those observed in vitro. For instance, while DNMT1 overexpression in mESCs does not affect embryoid body growth and morphology(44), a similar level of overexpression causes mid-gestation lethality in mouse embryos(51), comparable to DNMT1 deficiency. Therefore, developmental defects may still emerge despite transient epigenetic perturbations only occurring in early stages of development. Indeed, significant developmental delays and a wide range of morphological anomalies were observed in the majority of mid-gestation mouse embryos lacking the DNMT1o protein(240). DNMT1o is the oocyte-derived variant of DNMT1 that supports DNA methylation maintenance in mouse preimplantation embryos; DNMT1o deficiency permanently reduces DNA methylation levels in mouse embryos, especially at genomic imprints(241,242). DNMT1o-deficient mouse embryos thus exemplify transient epigenetic disruption during the early developmental stage of preimplantation, which precedes blastocyst formation. We previously demonstrated that DNMT1o deficiency during mouse embryonic development also impacts extraembryonic tissues, causing their hyperplasia(243), possibly due to the dispensability of DNA methylation for their growth and survival(191). Results from this study are consistent with those findings; the estimated proportion of extraembryonic endoderm cells was significantly higher in DNMT1-deficient mouse embryoid bodies compared to controls. In contrast, rescuing DNMT1 in mESCs before the formation of embryoid bodies prevented this outcome, though whether extraembryonic hyperplasia would be fully prevented in vivo remains an intriguing question. This could be explored by injecting the *Dnmt1*-rescue mESCs from this study into blastocyst-stage mouse embryos to gain deeper insights into how the repercussions of epigenetic perturbations may become intensified in the complex environment of embryonic development compared to in vitro models.

In conclusion, rescuing DNMT1 activity was insufficient to reverse the effects of its loss in mESCs; only partial reversal was achieved. Permanent effects were detected as well as effects triggered by the rescue intervention, highlighting the difficulty of completely reversing the impact of epigenetic perturbations without unintended consequences. The effects of transient loss of DNMT1 in mESCs extended well beyond DNA methylation alterations, encompassing broader molecular and cellular alterations. Disruption of histone modification landscapes and gene expression profiles were observed, along with transposable element de-repression, genomic instability, as well as morphological and functional alterations, underscoring the far-reaching influence of DNMT1. Transient loss of DNMT1 in mESCs also enabled the discovery 20 regions with imprinted-like epigenetic and regulatory signatures, some of which may be relevant to human health. Finally, we demonstrated that rescuing DNMT1 activity after differentiation onset exacerbated molecular and cellular disturbances, enlightening the heightened sensitivity of differentiating cells to epigenetic perturbations and the favourability of intervening during stages of greater cellular plasticity. These findings broaden the scope and complexity of DNMT1-dependent mechanisms, emphasize the wide-ranging repercussions of modulating key epigenetic enzymes and provide insight into the potential and challenges of therapeutic interventions aimed at reversing the effects of epigenetic perturbations during embryonic development.

## LIMITATIONS OF THE STUDY

This study presents some limitations that warrant consideration. Firstly, EM-seq does not distinguish 5-methylcytosine (5mC; methylation) from 5-hydroxymethylcytosine (5hmC; hydroxymethylation), an intermediate product of DNA demethylation by Ten-eleven translocation (TET) enzymes. However, given the low global 5hmC levels in mESCs (∼3%)(244) and the stable expression of *Tet1-3* in our model, it is unlikely that 5hmC substantially skewed our results. Moreover, lack of parental single nucleotide polymorphisms in *Dnmt1*^tet/tet^ mESCs prevented allele-specific profiling of epigenetic modifications and gene expression, thereby constraining the depth of our investigation into genomic imprinting maintenance. Additionally, male mESCs were chosen to avoid X inactivation effects and enable analysis of both sex chromosomes, therefore no sex-specific analyses were performed. However, considering that transient *Dnmt1* inactivation led to DNA hypomethylation of sex chromosomes in mESCs and dysregulation of X- and Y-linked genes in mESCs and upon their differentiation into germ layers, our understanding of sex-specific responses to epigenetic disruptions could be enhanced by additional experiments using a female *Dnmt1*^tet/tet^ mESC model. Finally, alternative methods for inducing transient loss of DNMT1 in WT mESCs were not attempted, for example via DNMT1 inhibitors (e.g. 5-azacytidine(245), decitabine(246), GSK-3484862(246,247)) or siRNA interference(248). Compared to the Tet-Off system, these methods often exhibit poor reproducibility across entire cell populations, with some inhibitors also lacking DNMT specificity and having cytotoxic properties. The *Dnmt1*^tet/tet^ model was therefore more suitable for a direct investigation of DNMT1-dependent mechanisms.

## AUTHOR CONTRIBUTIONS

Conceptualization & methodology: E.E., S.M., V.B-L., T.D., N.J-M.R., N.G., G.B., D.S.; Investigation: E.E., L-M.L.; Formal analysis: E.E., A.L., M.C.; Writing–original draft: E.E., S.M.; Writing–review & editing: All authors; Visualization: E.E.; Supervision: S.M., G.B., D.S.; Funding acquisition: S.M., N.G.

## DATA AND CODE AVAILABILITY

The Gene Expression Omnibus (GEO) accession number for raw and processed data produced in this study will be made available upon the publication of this paper. The GEO accession numbers for published data utilized in this study are GSE123942 (*Zfp57* WT and KO mESCs) and GSE171749 (*Setdb1* WT and KO mESCs). Allele-specific methylated regions in mESCs (Leung 2014) and CpG methylation profiles of mouse sperm and oocytes (Wang 2014) are accessible via the UCSC table browser from the DNA methylation Track Hub. Single-cell RNA sequencing data used for deconvolution of embryoid body mRNA sequencing raw counts were obtained from the R package MouseGastrulationData v1.18.0. This study does not report original code. However, scripts for data analysis can be provided upon request to lead contact.

## ACKNOWLEDGEMENTS

We thank the members of the McGraw laboratory for valuable insights and support throughout the course of this study, the McGill Genome Centre and *Génome Québec* for sequencing services, Ines Boufaied (Azrieli Research Centre of Sainte-Justine University Hospital) for assisting with FACS experiments, as well as Gregor Andelfinger (University of Montreal) and J. Richard Chaillet (University of Pittsburgh) for gifting R1 and *Dnmt1*^tet/tet^ mESCs, respectively.

