## Supplemental Figures for "Rescuing DNMT1 Fails to Fully Reverse the Molecular and Functional Repercussions of Its Loss in Mouse Embryonic Stem Cells"

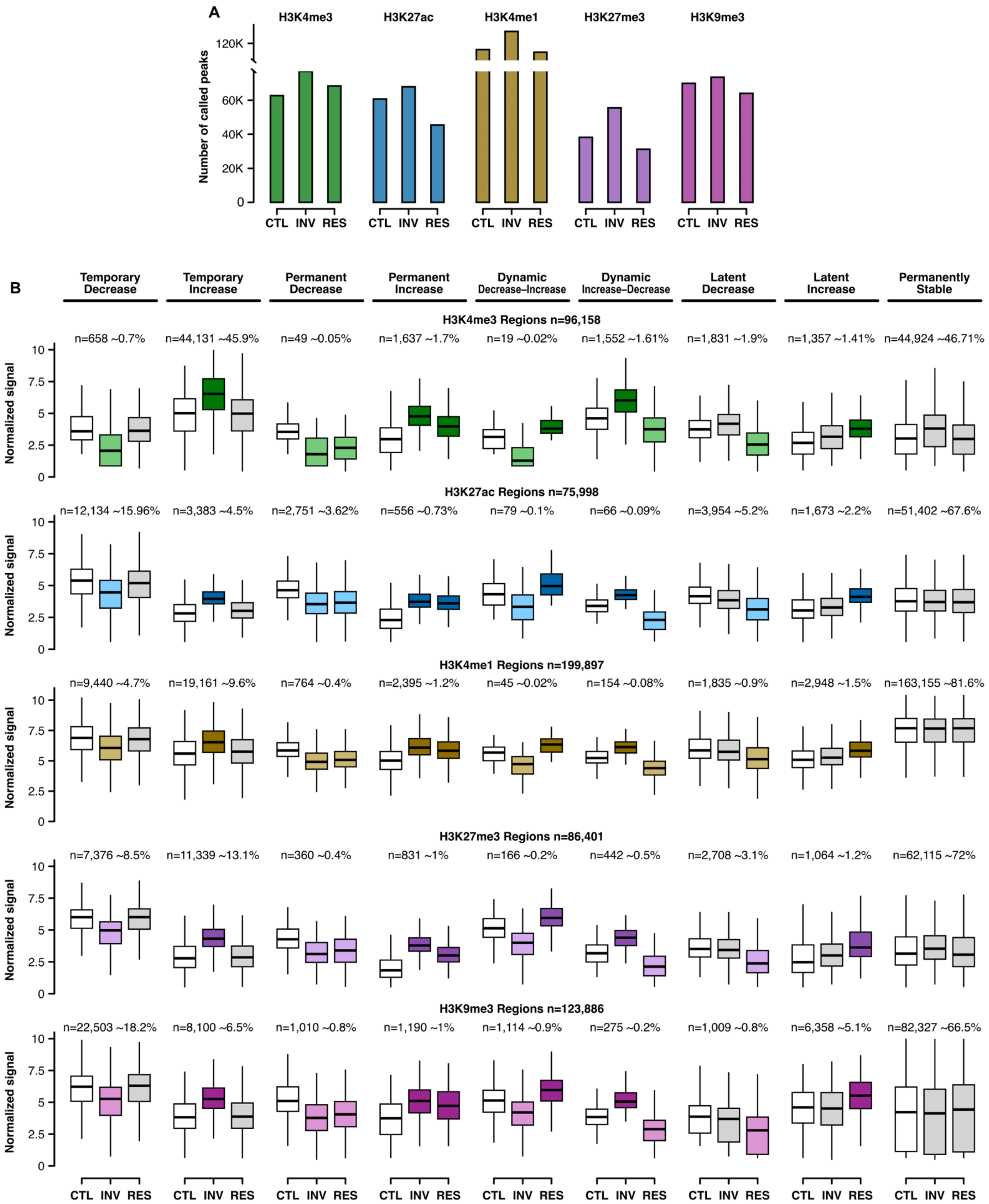


Boxplot outliers not shown

**Figure S1:** <u>Histone modification alterations during *Dnmt1* inactivation and rescue in mESCs.</u> **(A)** Number of called peaks for H3K4me3, H3K27ac, H3K4me1, H3K27me3 and H3K9me3 in *Dnmt1*$^{CTL}$, *Dnmt1*$^{INV}$ and *Dnmt1*$^{RES}$ using the MACS2 software. Histone modifications were measured by ChIP sequencing (n=2 per condition). **(B)** Alteration patterns of histone-modified regions based on differential states in *Dnmt1*$^{INV}$ and *Dnmt1*$^{RES}$ versus *Dnmt1*$^{CTL}$ identified in Figures 2A-B using the MAnorm2 software. Temporary: decrease/increase in *Dnmt1*$^{INV}$ & stable in *Dnmt1*$^{RES}$. Permanent: decrease/increase in both *Dnmt1*$^{INV}$ & *Dnmt1*$^{RES}$. Dynamic: decrease/increase in *Dnmt1*$^{INV}$ & the opposite in *Dnmt1*$^{RES}$. Latent: stable in *Dnmt1*$^{INV}$ & decrease/increase in *Dnmt1*$^{RES}$. *Related to Figure 2.*

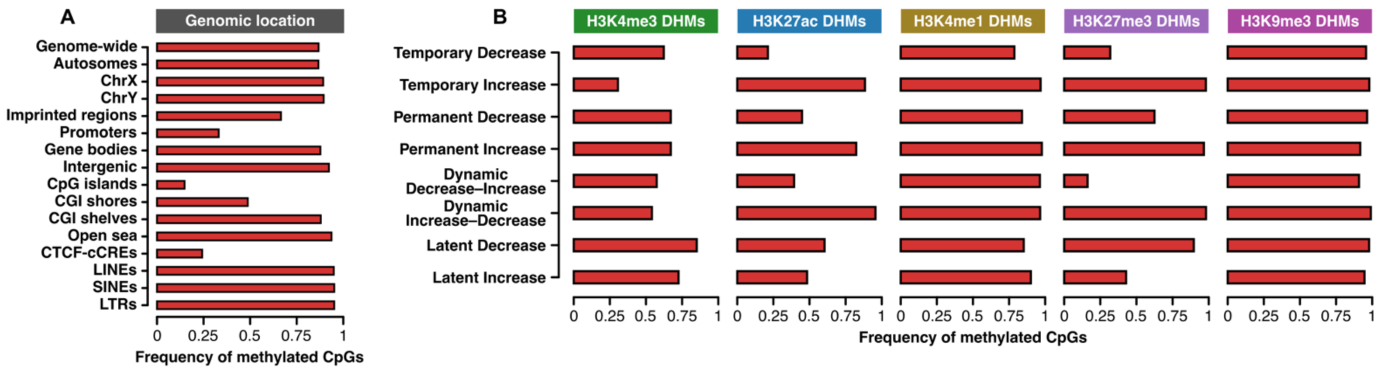


**Figure S2:** <u>CpG methylation in *Dnmt1*$^{CTL}$ mESCs relative to genomic location context and histone modification alteration patterns.</u> **(A)** Frequency of methylated CpGs in *Dnmt1*$^{CTL}$ relative to various genomic location contexts. **(B)** Frequency of methylated CpGs in *Dnmt1*$^{CTL}$ relative to differentially histone-modified regions (DHMs) categorized into alteration patterns identified in Figure 2C. Methylated CpGs were determined by a methylation level ≥ 20% in *Dnmt1*$^{CTL}$. Frequency of methylated CpGs represents the number of methylated CpGs divided by the number of sequenced CpGs. *See also Tables S1 and S2*. *Related to Figure 3*.

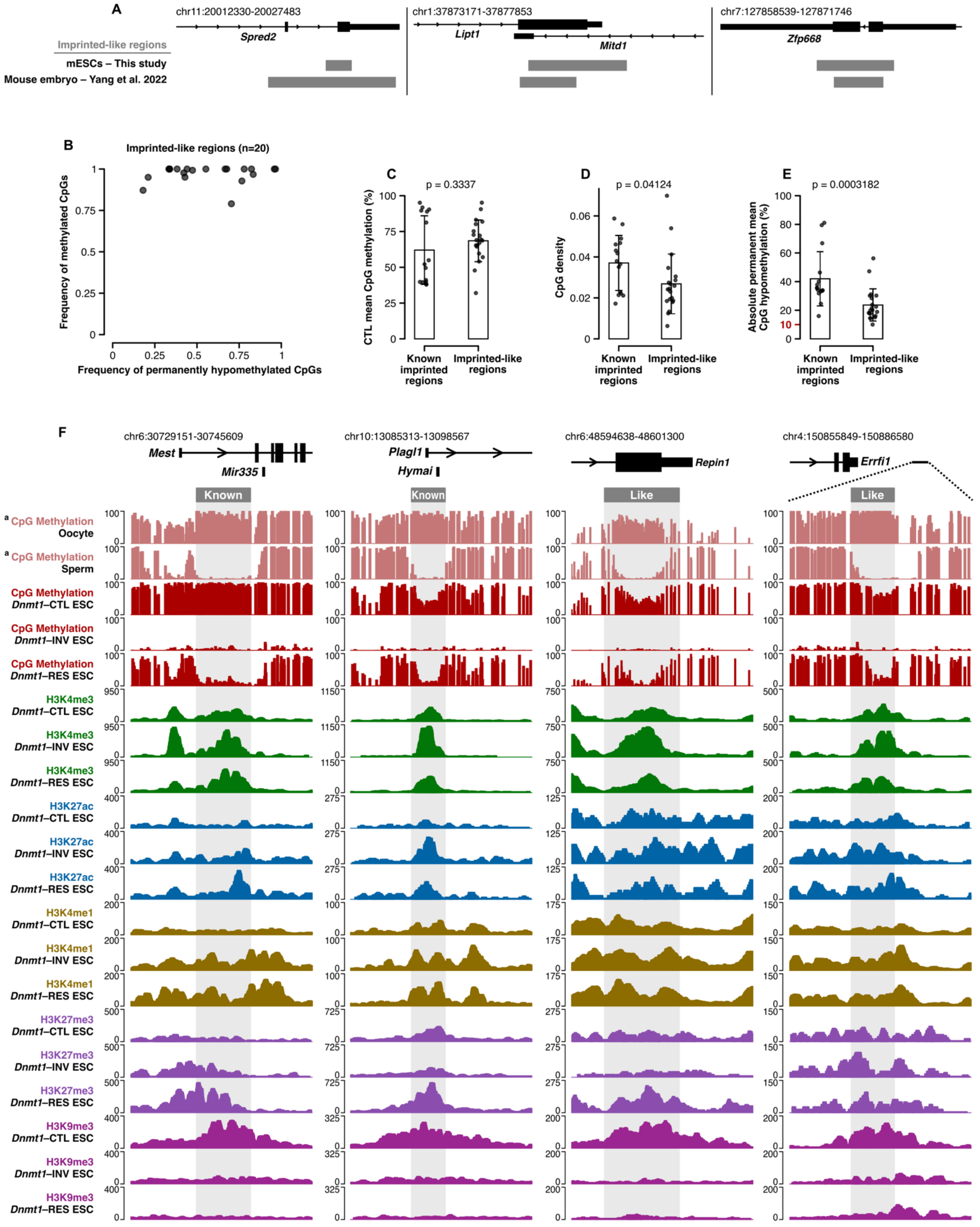


[a] Wang 2014 & Smith Lab

CpG methylation unit : percentage (%)    Histone modifications unit : Reads Per Kilobase per Million mapped reads (RPKM)

**<u>Figure S3:</u>** <u>Analysis of known imprinted regions enables the identification of 20 regions with imprinted-like epigenetic and regulatory signatures.</u> **(A)** Overlap of *Spred2*, *Lipt1-Mitd1* and *Zfp668* imprinted-like regions from our study with imprinted-like regions identified by Yang et al. 2022 in mouse preimplantation embryos. **(B)** Frequency of methylated CpGs versus frequency of permanently hypomethylated CpGs for imprinted-like regions identified in Figure 4B. **(C-E)** Comparison of mean CpG methylation levels in *Dnmt1*$^{CTL}$ (Kruskal-Wallis test), CpG density (Student's t test) and absolute permanent mean CpG hypomethylation (Kruskal-Wallis test) between known imprinted regions selected in Figure 4A and the imprinted-like regions identified in Figure 4B. **(F)** Genomic signal tracks for *Mest* and *Plagl1* imprinted regions and *Repin1* and *Errfi1 downstream* imprinted-like regions showing CpG methylation in gametes and *Dnmt1*$^{tet/tet}$ mESCs as well as H3K4me3, H3K27ac, H3K4me1, H3K27me3 and H3K9me3 in *Dnmt1*$^{tet/tet}$ mESCs. *See also Table S3*. *Related to Figure 4*.

### Imprinted genes regulated by selected known imprinted regions n=40

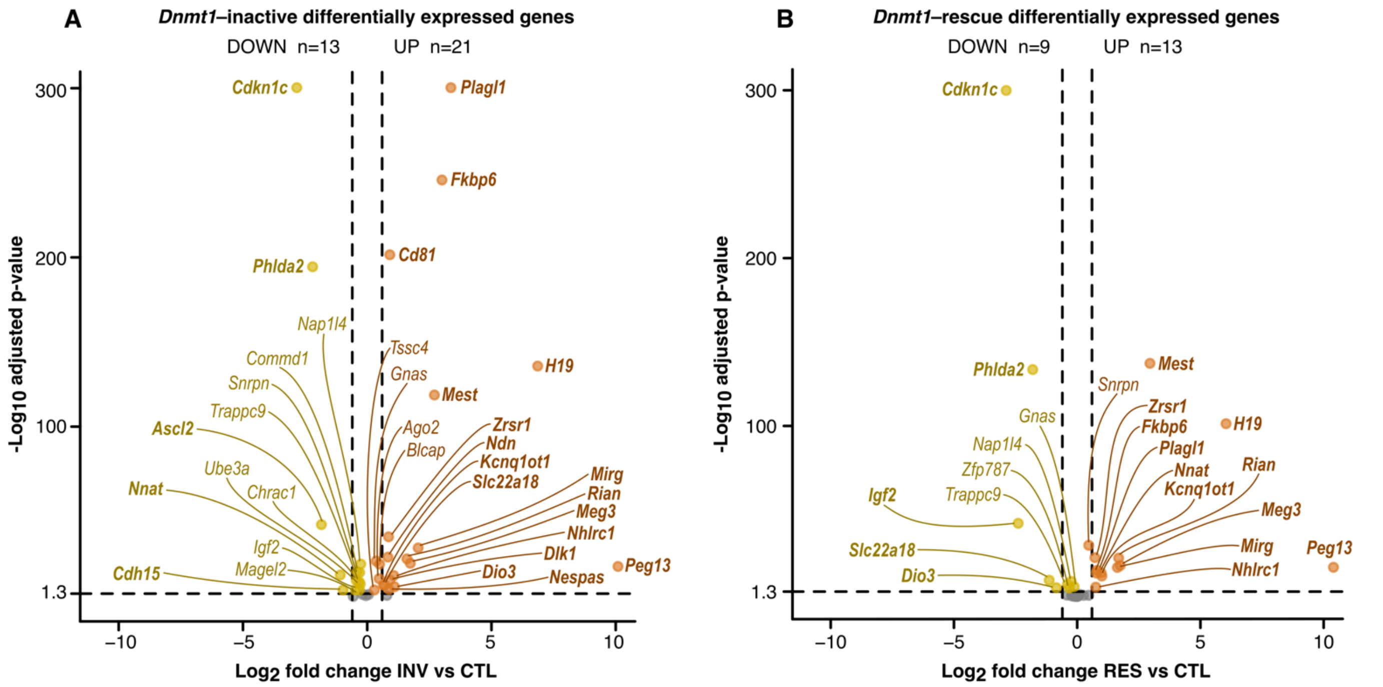


Stable INV vs CTL (n=6) : *Copg2, Mkrn3, Kcnq1, Zfp787, Rtl1, Kcnk9*

Stable RES vs CTL (n=18) : *Copg2, Magel2, Mkrn3, Ube3a, Chrac1, Ascl2, Commd1, Cdh15, Nespas, Kcnq1, Tssc4, Dlk1, Ndn, Ago2, Rtl1, Kcnk9, Blcap, Cd81*

Horizontal dashed lines : Adjusted p-value 0.05  Vertical dashed lines : $Log_2$ fold change -0.6 and 0.6

Bold gene names : $Log_2$ fold change ≤ -0.6 or ≥ 0.6

### Genes related to imprinted-like regions n=22

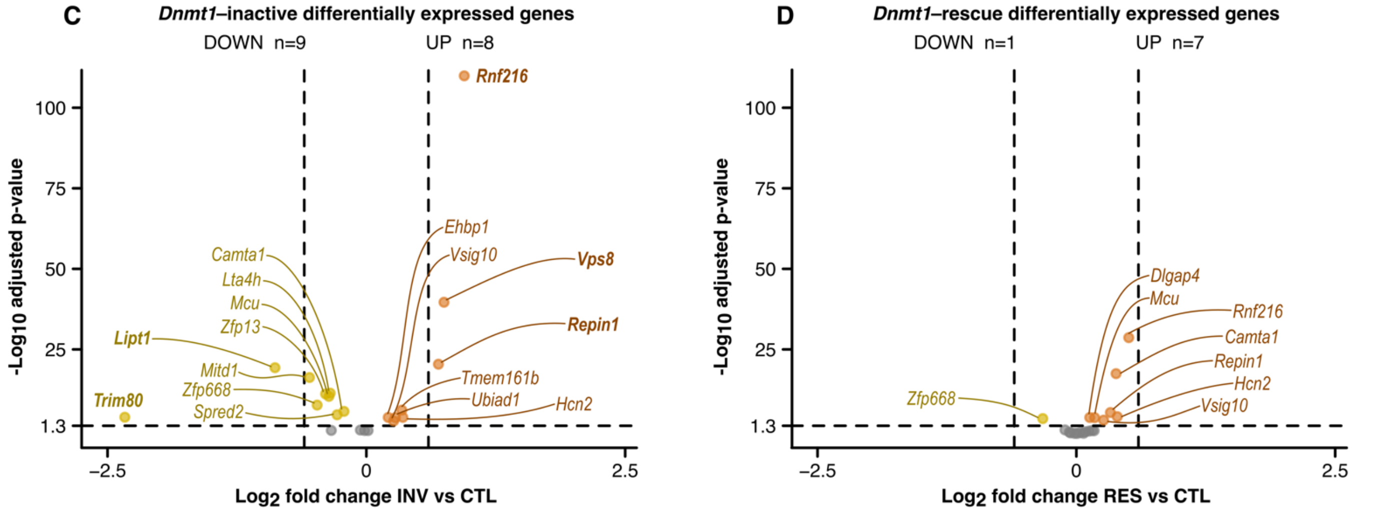


Stable INV vs CTL (n=5) : *Mir6378, Dlgap4, Polrmt, Gsc, Errfi1*

Stable RES vs CTL (n=14) : *Mir6378, Tmem161b, Ubiad1, Vps8, Ehbp1, Trim80, Lta4h, Spred2, Zfp13, Lipt1, Mitd1, Polrmt, Gsc, Errfi1*

Horizontal dashed lines : Adjusted p-value 0.05  Vertical dashed lines : $Log_2$ fold change -0.6 and 0.6

Bold gene names : $Log_2$ fold change ≤ -0.6 or ≥ 0.6

**Figure S4:** Differential expression of genes linked to known imprinted regions and imprinted-like regions. **(A-B)** Differential expression in *Dnmt1*$^{INV}$ and *Dnmt1*$^{RES}$ versus *Dnmt1*$^{CTL}$ of genes regulated by the 15 known imprinted regions selected in Figure 4A assessed by an adjusted p-value ≤ 0.05 using mRNA sequencing (n=3 per condition) analyzed with the DESeq2 software. **(C-D)** Differential expression in *Dnmt1*$^{INV}$ and *Dnmt1*$^{RES}$ versus *Dnmt1*$^{CTL}$, respectively, of genes associated with the imprinted-like regions identified in Figure 4B assessed by an adjusted p-value ≤ 0.05 using mRNA sequencing analyzed with the DESeq2 software. *See also Table S4*. *Related to Figure 4*.

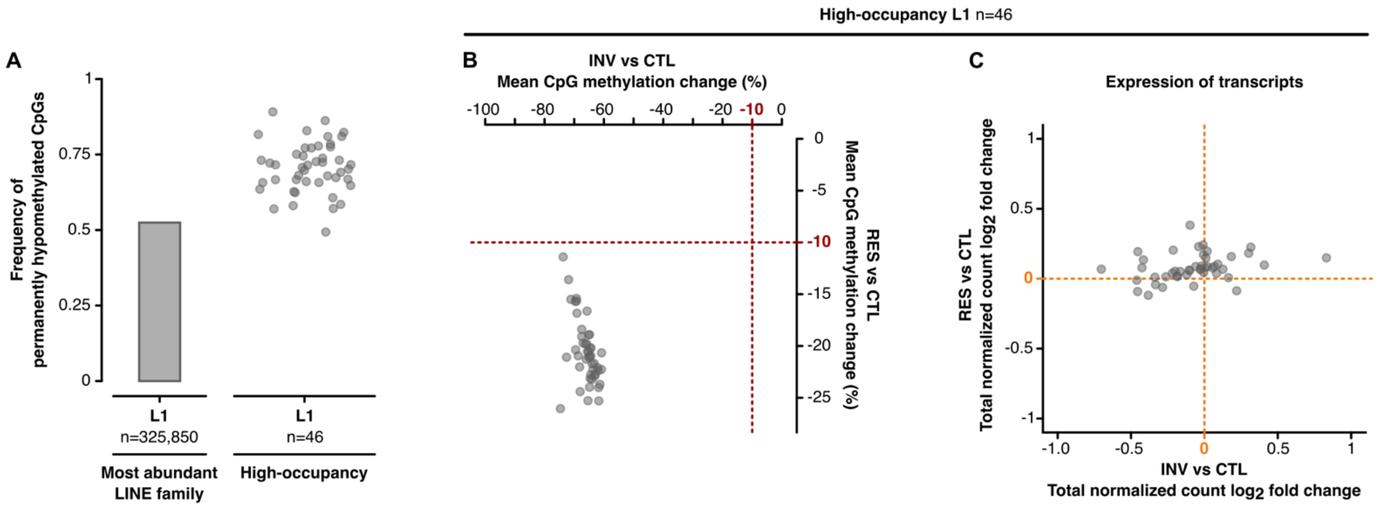


**Figure S5:** Permanent CpG hypomethylation of high-occupancy L1 LINEs throughout *Dnmt1* inactivation and rescue in mESCs is not linked to major changes in their transcription. **(A)** Frequency of permanently hypomethylated CpGs in L1 LINEs, the most abundant LINE family, and in high-occupancy L1 LINEs (genomic occurrences ≥ 500). **(B-C)** Change in mean CpG methylation levels (EM-seq) and $log_2$ fold change of total RNA-seq normalized counts of high-occupancy L1 LINEs in *Dnmt1*$^{INV}$ and *Dnmt1*$^{RES}$ versus *Dnmt1*$^{CTL}$ (EM-seq: n=3, RNA-seq: n=2 per condition).

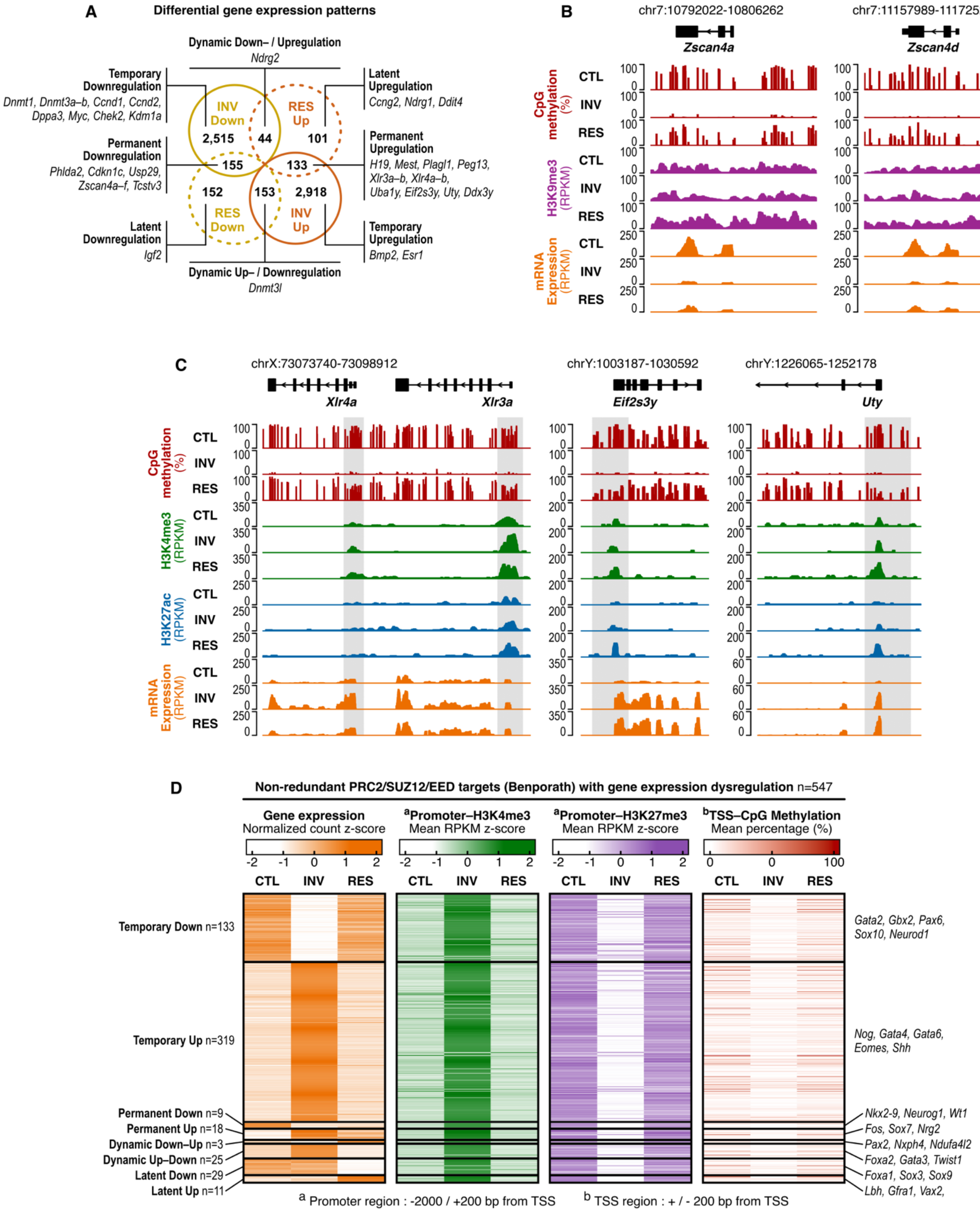
A
Differential gene expression patterns
Dynamic Down– / Upregulation
Ndrg2
Temporary Downregulation
Dnmt1, Dnmt3a–b, Ccnd1, Ccnd2, Dppa3, Myc, Chek2, Kdm1a
Latent Upregulation
Ccng2, Ndrg1, Ddit4
INV Down
RES Up
2,515
44
101
Permanent Downregulation
Phlda2, Cdkn1c, Usp29, Zscan4a–f, Tcstv3
155
133
Permanent Upregulation
H19, Mest, Plagl1, Peg13, Xlr3a–b, Xlr4a–b, Uba1y, Eif2s3y, Uty, Ddx3y
152
153
2,918
RES Down
INV Up
Latent Downregulation
Igf2
Temporary Upregulation
Bmp2, Esr1
Dynamic Up– / Downregulation
Dnmt3l
B
chr7:10792022-10806262
chr7:11157989-11172521
Zscan4a
Zscan4d
CpG methylation (%)
H3K9me3 (RPKM)
mRNA Expression (RPKM)
CTL
INV
RES
C
chrX:73073740-73098912
chrY:1003187-1030592
chrY:1226065-1252178
Xlr4a
Xlr3a
Eif2s3y
Uty
CpG methylation (%)
H3K4me3 (RPKM)
H3K27ac (RPKM)
mRNA Expression (RPKM)
D
Non-redundant PRC2/SUZ12/EED targets (Benporath) with gene expression dysregulation n=547
Gene expression
Normalized count z-score
aPromoter–H3K4me3
Mean RPKM z-score
aPromoter–H3K27me3
Mean RPKM z-score
bTSS–CpG Methylation
Mean percentage (%)
CTL INV RES
Temporary Down n=133
Temporary Up n=319
Permanent Down n=9
Permanent Up n=18
Dynamic Down–Up n=3
Dynamic Up–Down n=25
Latent Down n=29
Latent Up n=11
Gata2, Gbx2, Pax6, Sox10, Neurod1
Nog, Gata4, Gata6, Eomes, Shh
Nkx2-9, Neurog1, Wt1
Fos, Sox7, Nrg2
Pax2, Nxph4, Ndufa4l2
Foxa2, Gata3, Twist1
Foxa1, Sox3, Sox9
Lbh, Gfra1, Vax2,
a Promoter region : -2000 / +200 bp from TSS
b TSS region : + / - 200 bp from TSS

**Figure S6:** Gene expression dysregulation during inactivation and rescue of *Dnmt1* in mESCs is associated with alteration of epigenetic modifications. **(A)** Gene expression alteration patterns based on differential states in *Dnmt1*$^{INV}$ and *Dnmt1*$^{RES}$ versus *Dnmt1*$^{CTL}$ identified in Figures 6A and 6B. Temporary: down/up in *Dnmt1*$^{INV}$ & stable in *Dnmt1*$^{RES}$. Permanent: down/up in both *Dnmt1*$^{INV}$ & *Dnmt1*$^{RES}$. Dynamic: down/up in *Dnmt1*$^{INV}$ & the opposite in *Dnmt1*$^{RES}$. Latent: stable in *Dnmt1*$^{INV}$ & down/up in *Dnmt1*$^{RES}$. **(B)** Genomic signal tracks showing CpG methylation, H3K9me3 and mRNA expression for *Zscan4a* and *Zscan4d* in *Dnmt1*$^{CTL}$, *Dnmt1*$^{INV}$ and *Dnmt1*$^{RES}$. **(C)** Genomic signal tracks showing CpG methylation, H3K4me3, H3K27ac and mRNA expression for *Xlr4a*, *Xlr3a*, *Eif2s3y*, and *Uty* in *Dnmt1*$^{CTL}$, *Dnmt1*$^{INV}$ and *Dnmt1*$^{RES}$. **(D)** Z-score normalization of mRNA-seq normalized count, promoter H3K4me3 and H3K27me3 levels, and TSS region CpG methylation level for PRC2/SUZ12/EED non-redundant targets with gene expression dysregulation in *Dnmt1*$^{INV}$ and/or *Dnmt1*$^{RES}$ identified in Figure 6D, categorized based on gene expression alteration patterns. *See also Tables S6 and S7*. *Related to Figure 6*.

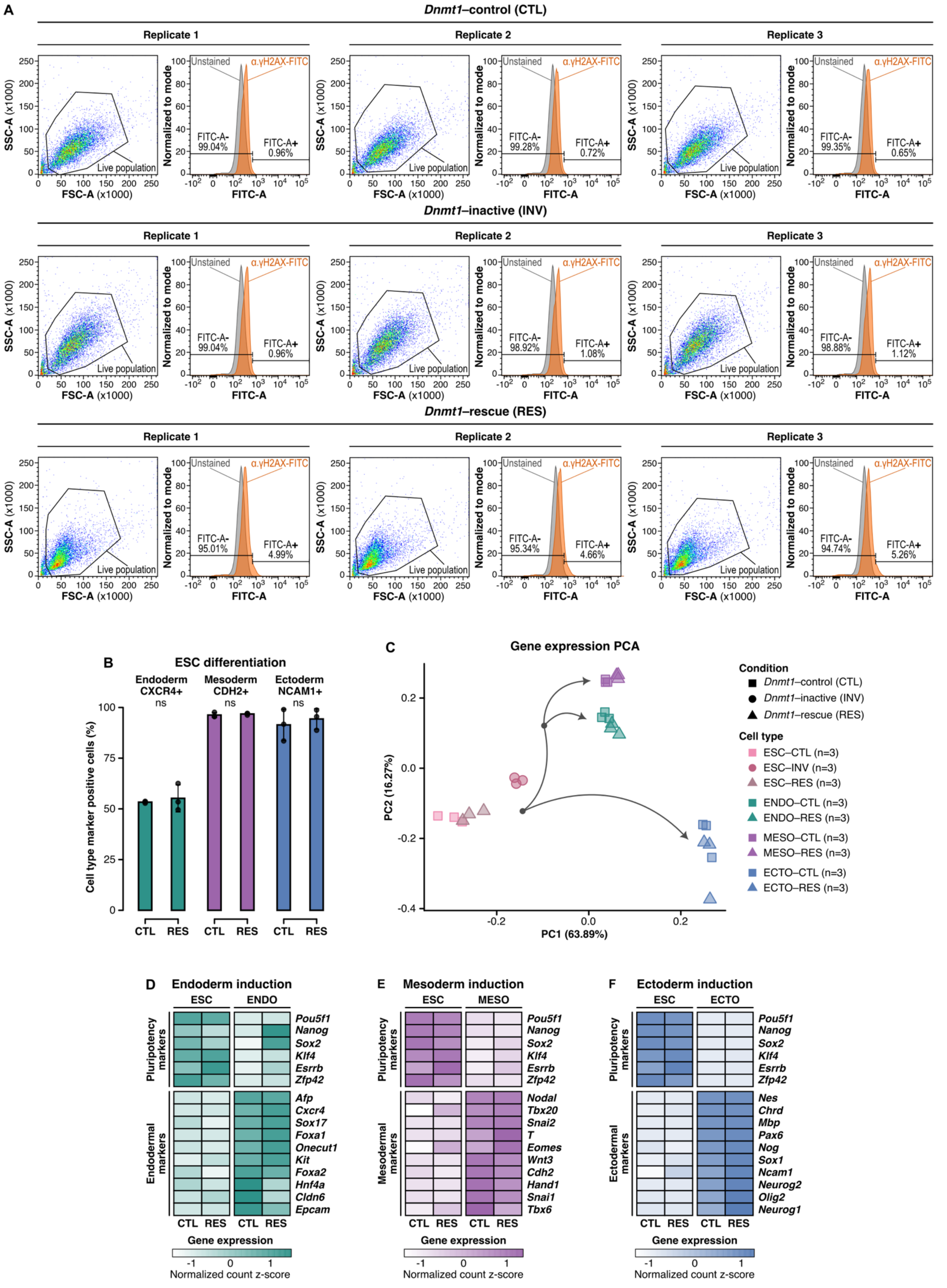

A
Dnmt1–control (CTL)
Dnmt1–inactive (INV)
Dnmt1–rescue (RES)
Replicate 1
Replicate 2
Replicate 3
SSC-A (x1000)
FSC-A (x1000)
Live population
Normalized to mode
Unstained
α.γH2AX-FITC
FITC-A
B
ESC differentiation
Endoderm CXCR4+
Mesoderm CDH2+
Ectoderm NCAM1+
ns
Cell type marker positive cells (%)
CTL
RES
C
Gene expression PCA
PC1 (63.89%)
PC2 (16.27%)
Condition
Dnmt1–control (CTL)
Dnmt1–inactive (INV)
Dnmt1–rescue (RES)
Cell type
ESC–CTL (n=3)
ESC–INV (n=3)
ESC–RES (n=3)
ENDO–CTL (n=3)
ENDO–RES (n=3)
MESO–CTL (n=3)
MESO–RES (n=3)
ECTO–CTL (n=3)
ECTO–RES (n=3)
D
Endoderm induction
E
Mesoderm induction
F
Ectoderm induction
ESC
ENDO
MESO
ECTO
Pluripotency markers
Endodermal markers
Mesodermal markers
Ectodermal markers
Gene expression
Normalized count z-score

**Figure S7:** Genomic instability and altered cellular processes due to inactivation and rescue of *Dnmt1* in mESCs. **(A)** FACS experiment detecting the presence of γH2AX, a DNA damage marker, in *Dnmt1^CTL^*, *Dnmt1^INV^* and *Dnmt1^RES^* live cell populations (n=3 per condition). Live cell populations were determined by forward scatter area (FSC-A) and side scatter area (SSC-A) measurements. FSC-A is an indication of cell size whereas SSC-A is an indication of cell granularity/internal complexity. **(B)** Percentage of viable endoderm, mesoderm and ectoderm cells derived from *Dnmt1^CTL^* and *Dnmt1^RES^* mESCs positive for their respective cell surface marker measured by FACS (n=3 per differentiation). Cell viability was assessed using propidium iodide staining. Percentage comparisons were conducted using Student's t test. **(C)** Principal component analysis of gene expression measured by mRNA sequencing in *Dnmt1^CTL^*, *Dnmt1^INV^* and *Dnmt1^RES^* mESCs and germ layers derived from *Dnmt1^CTL^* and *Dnmt1^RES^* mESCs (n=3 per cell type of each condition). Grey curved arrows show typical differentiation trajectories of mESCs towards germ layers. Endoderm and mesoderm first pass through a common mesendoderm stage, whereas the ectoderm follows a distinct differentiation trajectory from the other two germ layers. **(D-F)** Z-score normalization of mRNA-seq normalized counts for pluripotency and germ layer markers throughout differentiation inductions of *Dnmt1^CTL^* and *Dnmt1^RES^* mESCs into endoderm, mesoderm, and ectoderm, respectively. *Related to Figures 6 and 7*.

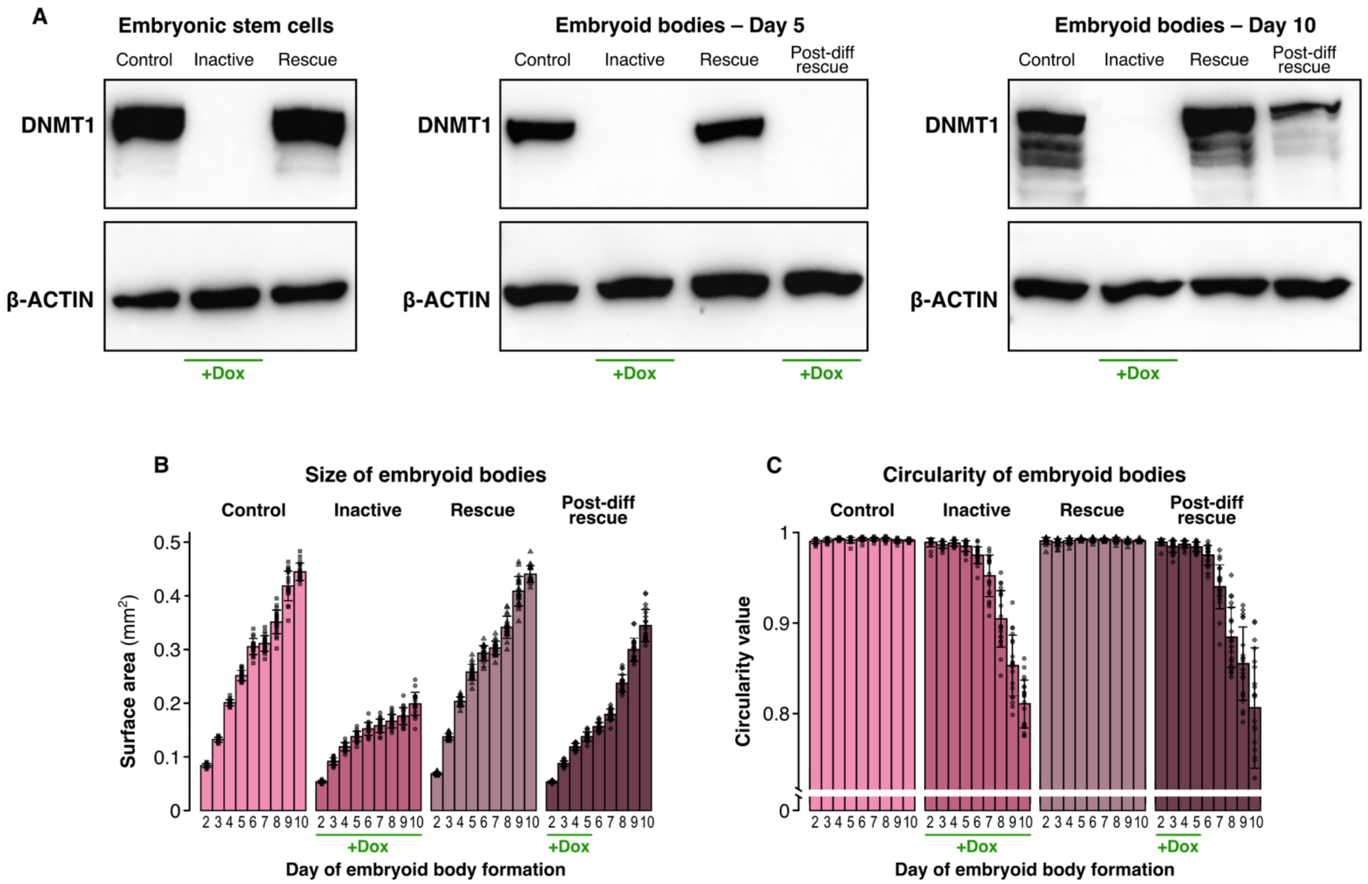


**Figure S8:** Delaying *Dnmt1* rescue until after differentiation initiation worsens molecular and cellular outcomes. **(A)** Western blot of DNMT1 in mESCs, day 5 embryoid bodies and day 10 embryoid bodies. **(B)** Size and **(C)** circularity of embryoid bodies from day 2 to day 10 determined by manually measuring their surface area and perimeter on bright-field microscopy images using the ImageJ software. Daily images were taken of the same 20 embryoid bodies per experimental condition. Surface area was used as an indicator of size and circularity was calculated using the formula 4π × Area/Perimeter². *Related to Figure 8*.
